# Intronic RNA of yeast *RPL22* paralogs acts as an allosteric switch

**DOI:** 10.64898/2026.02.10.704854

**Authors:** Kateřina Abrhámová, Alexandra Gredová, Karolína Navrátilová, Mohamed Boumaiza, Petr Folk

## Abstract

Ribosomal proteins, because of their RNA-binding capacity, may engage various cellular RNAs and fulfill nonribosomal roles. Previously, we and others described the intergenic regulation mediated by splicing of *RPL22* paralogs in *Saccharomyces cerevisiae.* Here, we prepared a panel of *RPL22A/B* intronic mutants with respect to their RNAfold-predicted features and analyzed their properties. We tested the splicing and Rpl22-intron interaction using an intron-containing reporter and a three-hybrid yeast system, respectively. We found that the splicing of *RPL22* introns can be inhibited by stabilizing a predicted stem as part of a particular type of conformation (I structure). Stabilizing the formation of an alternate stem (P structure) led to a permissive outcome of splicing. Intriguingly, the regulatory capacity of the main stem loop of the I structure was dependent on the rest of the intronic structure. Rpl22 enhanced splicing inhibition in WT and several of the mutants, which we interpret as stabilization of the I structure by protein binding. Mutagenesis identified both the main and alternative 5’ss and additional stem loops as part of the regulatory mechanism. The inhibitory conformation of the intron did not prevent recognition of the 5’ss and branch point, but rather stalled splicing at a later stage, before the first catalytic step. We concluded that the structural ensemble of the *RPL22* pre-mRNA behaves as an allosteric switch that responds to [Rpl22].

## Introduction

Ribosomal proteins (RP) constitute structural and regulatory components of ribosomes and as such remain highly conserved during evolution. Despite this constraint, some RPs acquired extra ribosomal functions by binding to RNAs other than rRNAs, mostly with an impact on gene expression [1]. As found, Rpl26 bound to the 5’untranslated region (5’UTR) of p53 mRNA, which stimulated translation and thus contributed to the regulation of the DNA damage response [2]. Rpl13a, upon IFN-γ activated phosphorylation, was released from ribosomes and translationally inhibited several groups of mRNAs as part of the GAIT system, which directed transcript selective translational control in myeloid cells [3,4]. The pre-mRNA molecules, their introns and untranslated regions in particular, can thus be viewed as evolving structures, which gained affinity for preexisting RPs [5].

Ribosomal protein coding genes (RPGs) often underwent duplication and exist as paralog pairs. In yeast, which experienced whole genome duplication followed by loss of duplicates, RPGs, in their majority, retained their paralogs. These gene pairs were implicated in splicing mediated intergenic regulation [6,7]. However, only a handful of such pairs were proven to regulate each other’s expression, for example, *RPS14*, *RPS9*, and *RPL22* [8–12]. Incorporation of two different paralogs into ribosomes can give rise to ribosome heterogeneity. Proteomic and transcriptomic analyzes found evidence for the existence of ribosomal subpopulations that differed in composition, interactomes, and perhaps mRNA specificity [8,13–18]. In yeast, the localization of *ASH1* mRNA translation into the emerging bud during yeast cell division, which prevents mating type switching, required specific RP paralogs [19].

Extraribosomal functions of the Rpl22 paralogs have been documented in yeast (see below), fruit fly, zebrafish, mice, and humans [20–25]. Human Rpl22 interacted with human telomerase RNA [26] and was also sequestered, apparently in competition, by the Epstein-Barr virus encoded RNAs in infected cells, possibly as part of the viral strategy to interfere with translation and growth [27,28]. More recently, Zhang and co-workers demonstrated that the zebrafish Rpl22 and Rpl22-Like1 controlled morphogenesis during gastrulation. The proteins acted antagonistically to modulate the splicing of the Smad2 pre-mRNA, presumably in cooperation with HNRNP-A1 [22]. In yeast, the pair Rpl22A/B was shown to be part of the oxidative stress response. Because *RPL22A* but not *RPL22B* mRNA is rich in UUG and because the translation efficiency of UUG rich transcripts increased under stress, the translation of *RPL22A* became more efficient, leading to a change in the Rpl22A/Rpl22B ratio [29]. Rpl22 was also shown to be required for the translation of *IME1* mRNA and meiotic induction in *S. cerevisiae*, as *rpl22*Δ cells were unable to translate the *IME1* transcript due to its atypical 5’UTR [30].

The yeast *RPL22A* and *RPL22B* genes form a feedback loop that modulates the ratio of paralogous transcripts in response to changing levels of Rpl22 proteins [11,12]. The regulatory loop formed by the *RPL22* pair couples the fluctuations in free Rpl22 proteins ([Rpl22A] + [Rpl22B]) to the ratio of their mRNAs ([*RPL22A*]/[*RPL22B*]). The shifts in mRNA ratio were related to oxidative stress (see above) or disbalances in the free RP concentration caused by exposures to MMS or Cd [11], supporting the interpretation that tuning the isoform ratio helps cells react to changing environment. The cellular processes that distinguish the difference in 19 amino acids between Rpl22A and Rpl22B remain unknown; the effects on splicing are apparently mediated by both A and B proteins [12]. Information on the interface between the intron and Rpl22, and between this complex and the spliceosome, is also limited. Gabunilas and Chanfreau postulated the intronic region between nt 153 and 225 (positions here and throughout the text are numbered relative to the first nucleotide of the *RPL22B* intron) as ‘regulatory element’ and found that it was responsible for protein binding (Fig 3C, D in [11]). This is similar to our conclusions based on yeast three-hybrid assays (nt 165 to 236; see I2 in [12]). They also showed that a predicted stem formation between nt 182-188 and 214-220 is necessary for Rpl22-mediated regulation and can rescue splicing inhibition *in vitro* when added *in trans* as part of free RNA (nt 115 to 255 in [11]).

RNA forms an ensemble of interchanging conformations, some of which may form with high probability and have a functional impact, as in riboswitches or snRNAs. The prevalence of particular conformations can be tuned by biological regulators or environmental conditions such as ions or temperature [31]. The RNA structure content changes with cultivation conditions and stress, suggesting that the structural aspect is relevant for our understanding of gene expression [22,32,33]. RNA stems can mediate interactions over longer distances in transcripts, which impacts the outcome of splicing [34,35]. The splicing sequences of the 5’splice site (5’ss), branch point (BP) or 3’splice site (3’ss) were incorporated into the stems and thus blocked [36], or brought into proximity and strengthened [35,37,38]. Exons were looped out [39] or selected for alternative splicing through base-pairing interactions, such as in the case of multi-cluster mutually exclusive exons [40–43]. RNA binding proteins that modulated RNA structures thus affected the outcome of alternative splicing [44,45]. In addition to directly engaging functional sequences *in cis* or obliterating the access of regulatory RNA binding proteins, RNA structures can intercept the functioning of large RNP complexes operating in gene expression [46–51].

Here, we aimed to characterize the intron-Rpl22 interaction in more detail, using constructs for *in vivo* measurements of splicing efficiency and yeast three-hybrid system. We were able to confirm the findings originally reported by Gabunilas and Chanfreau on the importance of a stem loop structure within the *RPL22B* intron [11]. Mutations that lowered the structure’s stability abolished inhibition mediated by Rpl22. On the contrary, we were unable to confirm sequence specificity with respect to Rpl22 binding or splicing inhibition within this region, which they reported. Importantly, we showed that inhibition of *RPL22B* intron (*RPL22B*i) splicing can be caused by an RNA structure without Rpl22, provided that the structure is stabilized *in cis*. As the pre-mRNA can be predicted to exist in alternate conformations, we used site-directed mutagenesis to show that only one alternative base pairing arrangement mediates the inhibitory effect. Furthermore, we showed that the inhibitory mechanism requires both the 5’ss proximal and the BP proximal parts of the pre-mRNA and that it does not operate through splice site obliteration, but rather by stalling spliceosome assembly prior to the first catalytic step.

## Results

### *The RPL22* introns are predicted to form alternative structures

Previous reports found that a stem loop between nucleotides 153 and 246 of *RPL22B*i was involved in Rpl22 binding and splicing inhibition (see Introduction; [11,12]. We examined the sequence alignments of both *RPL22* paralogs together with the related yeast species of the Saccharomycotina group. We found that the introns share conserved sequence elements, which could base pair in alternate arrangements (Fig 1A). Only one of the two arrangements would give rise to the stem loop between nucleotides 153 and 246 of *RPL22B*i.

**Fig 1.**
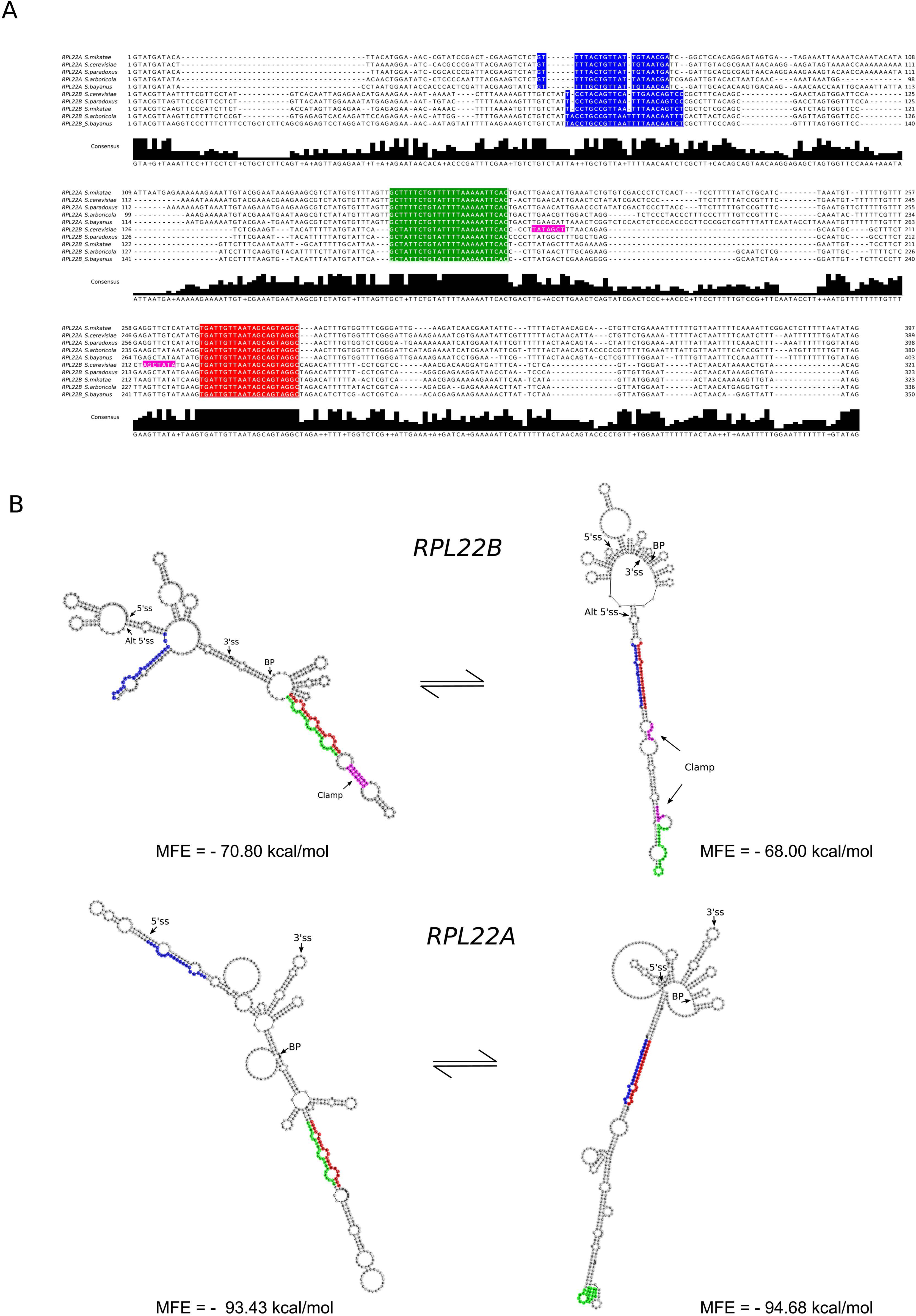
The *RPL22* introns are predicted to assume alternate conformations. **(A) The *RPL22* introns of Saccharomycotina yeasts contain complementary regions that suggest the formation of alternate secondary structures.** *The RPL22* introns of Saccharomycotina contain an absolutely conserved region of 23 nt positioned 5’ of the branch point (red), which can alternately base pair with partially complementary sequence elements located further upstream (green and blue). The homologs of *RPL22A* and *RPL22B* are grouped in the upper and lower rows of the alignment. Additional regions of complementarity are specific for the B paralogs (violet). The alignment was created using the MUSCLE algorithm https://www.ebi.ac.uk/Tools/msa/muscle/ and Jalview ver 2.11.2.6 was used for visualization. **(B) Predicted structural ensembles of intronic RNAs of *Saccharomyces cerevisiae RPL22A* and *RPL22B* paralogs contain two major types of stem loop arrangements.** Alternate structures of *RPL22* introns were predicted by RNAfold [52]. Pairing the conserved region (red in (A)) with the 3’complementary sequence element (3’COMP; green in (A)) leads to the formation of an inhibitory structure (denoted ‘I’), while the participation of the 5’complementary sequence element (5’COMP; blue in (A)) results in permissive structures (denoted ‘P’). Both structure types are predicted to occur in many variations and regardless of the inclusion of intron-bordering regions. The sequence range used for the modeling included 56 nt in front of the intron and 41 nt from exon 2; Minimum free energy (MFE) structures were visualized by RNAfold web server [52,53] and MFE values were calculated by RNAShapes Studio [54]. The colors correspond to the boxes in (A). The additional base pair complementarity in *RPL22B* (violet) is denoted as “Clamp”.

We revisited the structural predictions of the introns plus their surrounding pre-mRNA sequences employing RNAfold, Mfold [52,55], RNAShapes [56] and Sfold [57]. Consistently, the introns were predicted to give rise to structures with two alternative base pairing arrangements (see Fig 1A). A conserved sequence element (CONS; labeled red) paired with (i)a proximal upstream element (3’ complementary element (3’COMP); green) or alternatively, (ii) a distal upstream element (5’ complementary element (5’COMP); blue). The conserved element was identical in the Saccharomycotina group, while its complementary elements showed more variability. In *RPL22B*i, an additional short stem was predicted (Clamp; violet). We annotated the alternate structure types as inhibitory (I; Fig 1B left) and permissive (P; Fig 1B right), because we found that they differ with respect to splicing inhibition (see below). The sample conformations of these two types for both *RPL22A*i and *RPL22B*i are presented in Fig 1B as RNAfold predicted minimum free energy (MFE) structures; they have similar base pair content and thus similar relative stability (see the MFE values in Fig 1B). The I ⇋ P conversion would require the change of well above 20 Watson-Crick base pairs (S1 Fig), suggesting that an energy barrier may kinetically limit the transitions (see the Discussion).

The introns of the *RPL22* paralogs differ in sequence and length but share the propensity to form alternate stems 3’COMP/CONS (I) and 5’COMP/CONS (P). However, the proportion of the two types of structure in the structural ensembles of *RPL22B* is shifted to I compared to *RPL22A* (see S6 Fig for Sfold prediction). The two-dimensional plot of base pairing probabilities in the Boltzmann weighted structural ensemble of *RPL22B* shows that the stem loop between nucleotides 153 and 246 forms prominently (S6A Fig), while in the plot for *RPL22A*, both I and P signature densities are present (S6F Fig).

### *RPL22*i splicing can be differentially manipulated by stabilizing either I or P types of structure

The Rpl22-binding stem (I2 RNA fragment; [12]) does not involve 5’ss, BP, or 3’ss themselves. This led us to ask what the mechanism of splicing inhibition is. Using site-directed mutagenesis, we stabilized either of the predicted conformations (see above) and tested the RNA performance. We took advantage of our previous measurements of the effects of Rpl22 proteins on intron splicing of *RPL22B* and *RPL22A in vivo* [12].

In our reporter assays, we used the *rpl22a*Δ *rpl22b*Δ strain that harbors an empty or an Rpl22 producing pVTU260 plasmid. The reporter construct included the *RPL22B* (or other) intron surrounded by its own 5’untranslated region (5’UTR) and exon 1 and 60 nt of exon 2 fused in frame with the *CUP1* reporter at the 5’ and 3’ ends of the intron, respectively (see S9A Fig). Thus, the structural and functional characteristics of the pre-mRNA molecule between nt -56 and +381 (positions are numbered relative to the first nucleotide of the *RPL22B* intron) were preserved, including the alternative 5’splice site (ALT5’ss; 65-GTTTGT). Therefore, we could test the splicing of *RPL22B* in the absence or presence of Rpl22A and directly compare the reads (the protein is not essential in *S. cerevisiae*). We used only the expression of Rpl22A in our assays, as we previously demonstrated that either of the paralogous proteins inhibits splicing to the same extent [12]. To complement the reporter splicing assays, we used the three-hybrid system to test the binding of Rpl22A to the I2 fragment of the *RPL22Bi* mutants generated in this study. The outcomes are summarized in Figs. 2B to 5B. The details of spot tests and RNAfold prediction of the stem structures are documented in S7 Fig.

**Fig 2.**
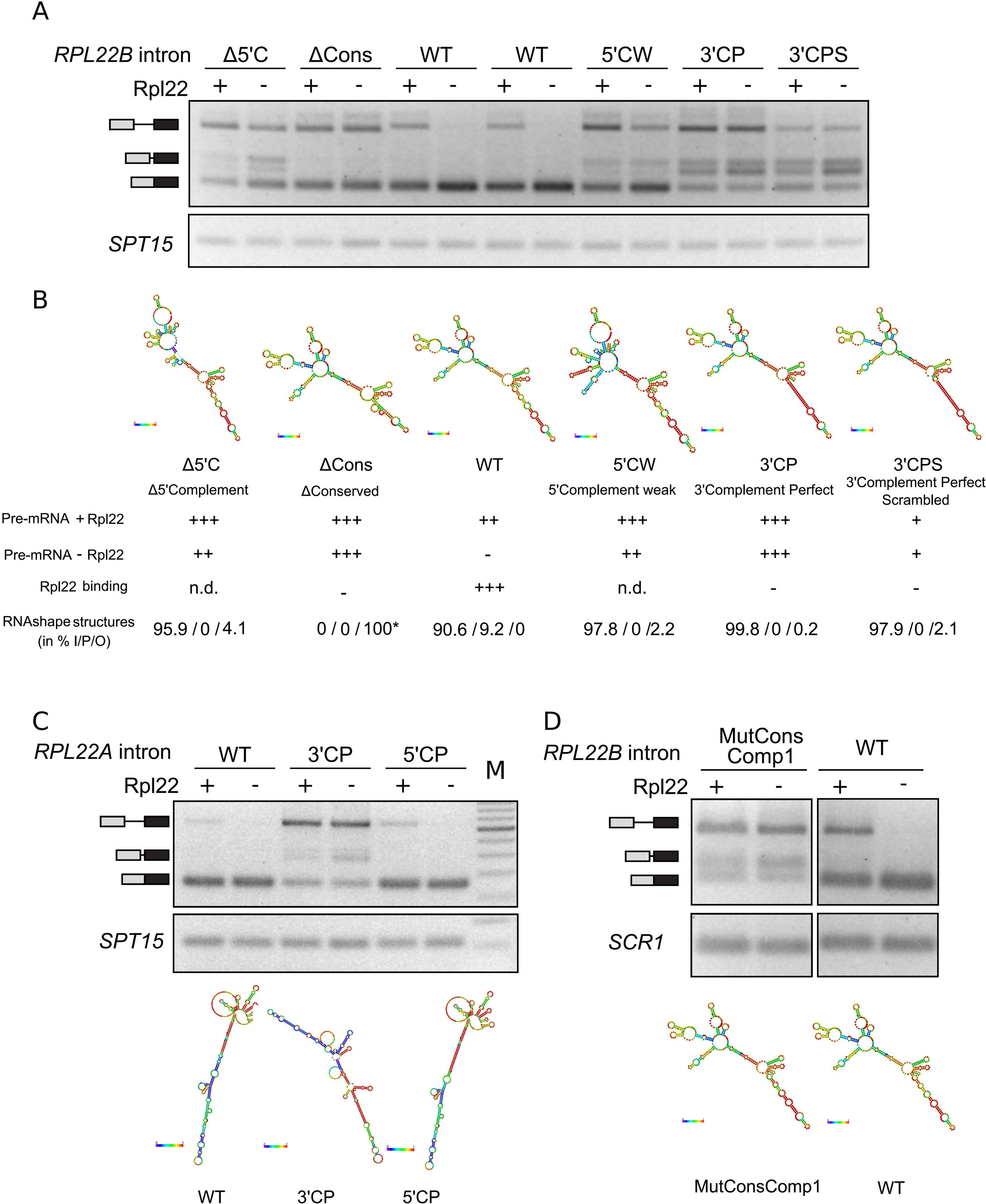
Splicing-inhibitory mutations block *RPL22* intron reporter splicing *in vivo* irrespective of the presence of the Rpl22 protein. **(A) Mutants stabilizing the inhibitory conformation of *RPL22B*i inhibited splicing irrespective of the presence of the Rpl22 protein *in vivo.*** The splicing efficiency of *RPL22B-CUP1* reporters was tested in *rpl22a*Δ *rpl22b*Δ strain harboring the plasmid pVTU260/RPL22A for overexpression of Rpl22A and in the same strain transformed with empty vector. Semi-quantitative PCR was run on cDNA prepared from RNA isolated from exponentially growing cultures using random hexamers. The loading control is included in a separate inset with the indicated gene. One of at least three independent experiments is shown. We detected alternative splicing products, which migrated above the size of the major product and which reflect the usage of alternative 5’ss. The PCR products of the corresponding length were sequenced to confirm this assumption. **(B) Summary of *RPL22B*i reporter experiments in (A), Rpl22 binding data from the yeast three-hybrid assay, and types of stem loop arrangements predicted by RNAshapes.** Pre-mRNA accumulation was approximated on a scale of ‘++ +’ to ‘-’. The binding of Rpl22 to a fragment of the *RPL22B* intron (I2 construct; see text) was tested in a yeast three-hybrid assay. Cell growth corresponding to Rpl22 binding was summarized on a scale from ‘++ +’ to ‘-’. For the photographs of spot tests, see S4 Fig; n.d. - “not done”. The secondary structures were predicted using RNAshapes [54]. The structures were sorted according to the stem loop arrangement involving the conserved region and its complementary elements (see Fig 1), and the number of structure types classified as I, P, or O (other) was expressed in %. See text and Figs. S1-S3 for additional information. (*) The 3’COMP/CONS base pairing in ΔCons was *sensu stricto* not of the I type, but the apical part of the main stem loop and the rest of the prediction remained the same as in the I structures (see also S2 Fig). **(C) The perfect complementarity between the conserved element and the 5’sequence element of *RPL22A*i inhibited the splicing similarly to *RPL22B*i.** The elements are annotated in Fig 1A. The experiment was carried out as in (A), except that the *RPL22A-CUP1* reporter was used. **(D) Keeping partial complementarity was sufficient to inhibit the splicing of *RPL22B*i with scrambled conserved element.** The mutant MutConsComp1 did not bind Rpl22 in Y3H (S7C Fig) and returned WT-like secondary structure prediction (S2 Fig). The experiment was done as in (A).

We reasoned that if the RNAfold predictions identified features that were relevant for the splicing inhibitory mechanism, their targeted disruption or hyperstabilization would promote or inhibit splicing, respectively. Indeed, we found that most of the manipulations that stabilized the 3’COMP/CONS base pairing (characteristic of I) had inhibitory and Rpl22 independent effects on splicing (Figs. 2 and S2). On the contrary, those mutations that destabilized the 3’COMP/CONS arrangement or favored the 5’COMP/CONS base pairing (characteristic of P) made the intron splicing permissive regardless of the presence of Rpl22A (Figs. 3 and S3). First, we stabilized the 3’COMP/CONS base pairing. In the 3’CP mutant, we made the 3’COMP element perfectly complementary to CONS; in 3’CPS, we kept the complementarity of the base pairs, but scrambled the base pairs; in the ΔCons, we deleted the whole CONS signature. These mutants had their presumed Rpl22 binding region mutated and were confirmed in Y3H assays as non-binding (Fig 2B). 3’CP and 3’CPS were predicted to form mostly the I structures (Figs. 2B and S2). In the splicing assay, we observed pre-mRNA accumulation with or without Rpl22A expressed with part of the signal recovered in bands that corresponded to ALT5’ss usage (Fig 2A). The deletion of CONS led to a structure that kept the key characteristics of the inhibitory mechanism intact. Although the base pairing of 3’COMP was *sensu stricto* not of the I type, the apical part, the main stem loop and the rest of the prediction remained the same (Figs 2 and S2). In addition to the mutant 3’CPS, which had a perfectly complementary main stem, we also designed the mutant MutConsComp1, which had a scrambled CONS motif in a stem of imperfect complementarity. The prediction of its secondary structure was close to WT (Figs 2 and S2). While the binding of Rpl22 was undetectable in Y3H (S7 Fig), the pre-mRNA structure retained its inhibitory function (Fig 2D).

**Fig 3.**
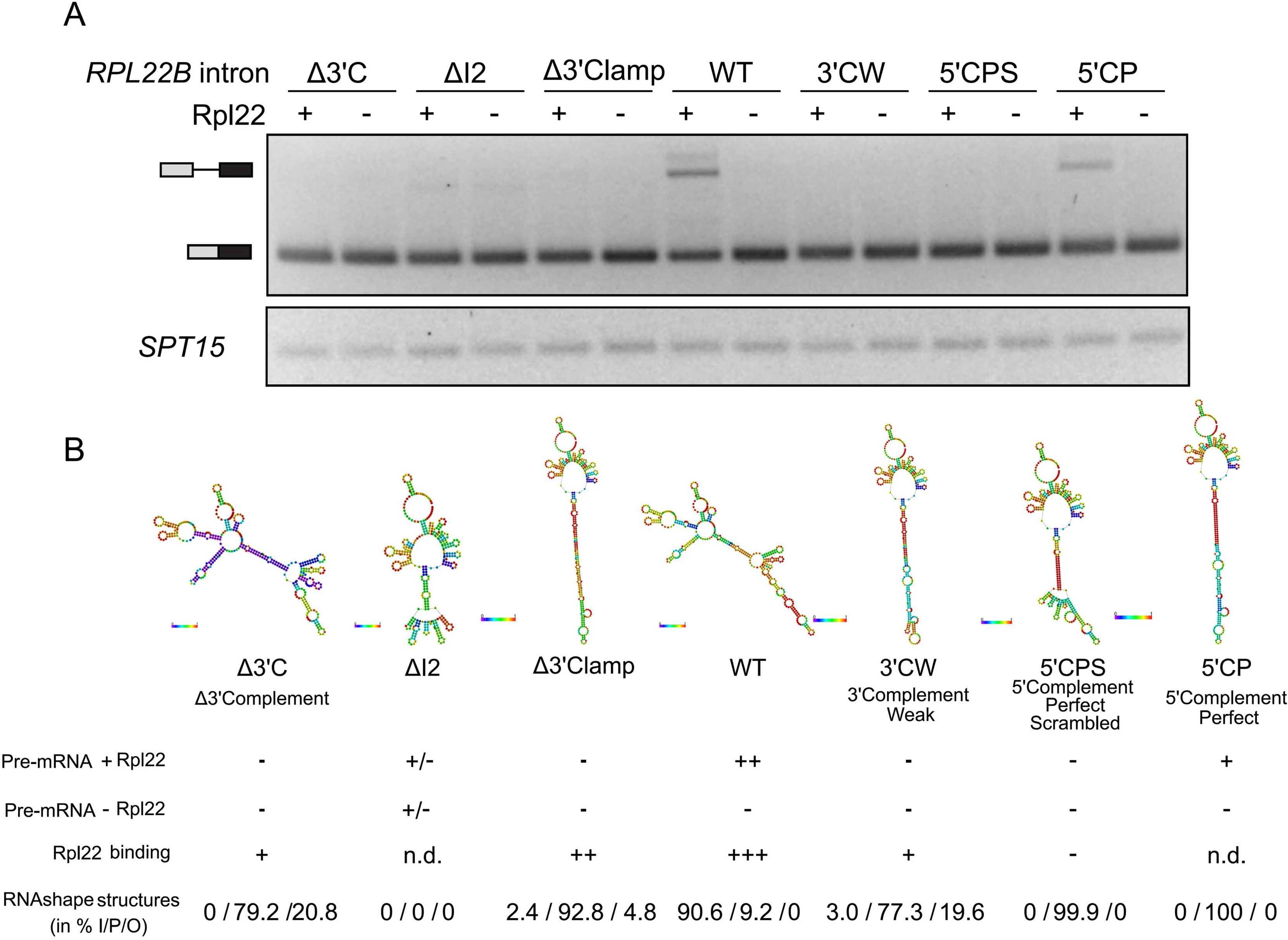
Splicing-permissive mutations did not block *RPL22B*i splicing *in vivo* irrespective of the presence of the Rpl22 protein. The experiment was done and the figure was formatted as in Fig 2. **(A) Mutants disrupting the inhibitory conformation of *RPL22B*i were splicing permissive.** See the legend of Fig 2A. **(B) Summary of the reporter experiments in (A), Rpl22 binding data from the yeast three-hybrid assay, and types of stem loop arrangements predicted by RNAshapes.** See the legend of Fig 2B.

When we removed the 5’COMP element or weakened its base pairing with CONS, we also obtained inhibitory phenotypes, although partly sensitive to Rpl22A. Further manipulations showed that the constructs that did not form the main stem loop spliced well and Rpl22A did not inhibit splicing. They included the deletion of the 3’arm of the clamp stem (Δ3’Clamp), the truncation of the main stem (ΔI2), and the deletion of 3’COMP (Δ3’C). Symmetrically to the 3’COMP manipulations, the perfect complementarity between 5’COMP and CONS (predicted as P) led to a permissive outcome, although part of the inhibitory effect of Rpl22A was still retained. When the base pairs of the 5’COMP/CONS stem were scrambled, the structure was splicing permissive and Rpl22A-refractory. Most of the manipulations in Fig 3 were predicted to form P structures (Figs. 3B and S3). In the case of ΔI2, the structural stability of the remainder of the intron was lowered (S3 Fig - upper right). We concluded that the stem structure in the *RPLB22*i was necessary for down-regulation of the splicing of *RPL22B.* Favoring its formation led to splicing inhibition even without Rpl22A binding (see 3’CP and 3’CPS; Fig 2B), while destabilization of the stem relieved inhibition. The expression of Rpl22A increased splicing inhibition in WT, Δ5’C, Δ5’CW and 5’CP, which we interpret as stabilization of the inhibitory conformation by binding of the protein rather than a ‘direct inhibition’ due to obstruction of the splice sites or BP.

The effects of the 3’CP and 5’CP mutations were also demonstrated for the *RPL22A* intron (Fig 2C). Stabilization of the 3’COMP/CONS base pairing inhibited splicing irrespective of the presence of Rpl22A (3’CP). The stabilization of the 5’COMP/CONS stem (5’CP) did not affect the result relative to WT (= Rpl22A still inhibited splicing), probably because the 5’COMP/CONS base pairing in WT is already relatively strong. The 5’CP mutation had a less prominent impact in *RPL22B* as well, likely because the P structures are not bound by the protein. The *Rpl22A*i structures thus showed the same propensity as *RPL22B*i, despite the sequence differences between the two introns.

### The main stem loop of the inhibitory structure of *RPL22B*i is necessary but not sufficient for the inhibition of splicing

The inhibitory structure of *RPL22B*i (see Fig 1B top left and S3 Fig) contained stem loops predicted in the vicinity of the branch point in the 5’ direction. We found that mutations in this region had strikingly distinct effects on the inhibitory mechanism. The destabilization of the BP-proximal hairpin left the inhibition intact, but melting of the distal hairpin (distal from BP in the 5’ direction) rendered the structure splicing permissive (Fig 4). This phenotype was retained even in a mutant with stabilized main stem loop (3’CP+BPmut4). In contrast, complete ablation of the distal hairpin (dLoop2) or disruption of the stem proximal to BP in the 3’ direction (Mutclamp2) produced a WT phenotype. In a mutant that combined stabilization of the main stem loop with distal hairpin deletion (3’CP+dLoop2), the structure still inhibited splicing, although inhibition was slightly enhanced by Rpl22A expression.

**Fig 4.**
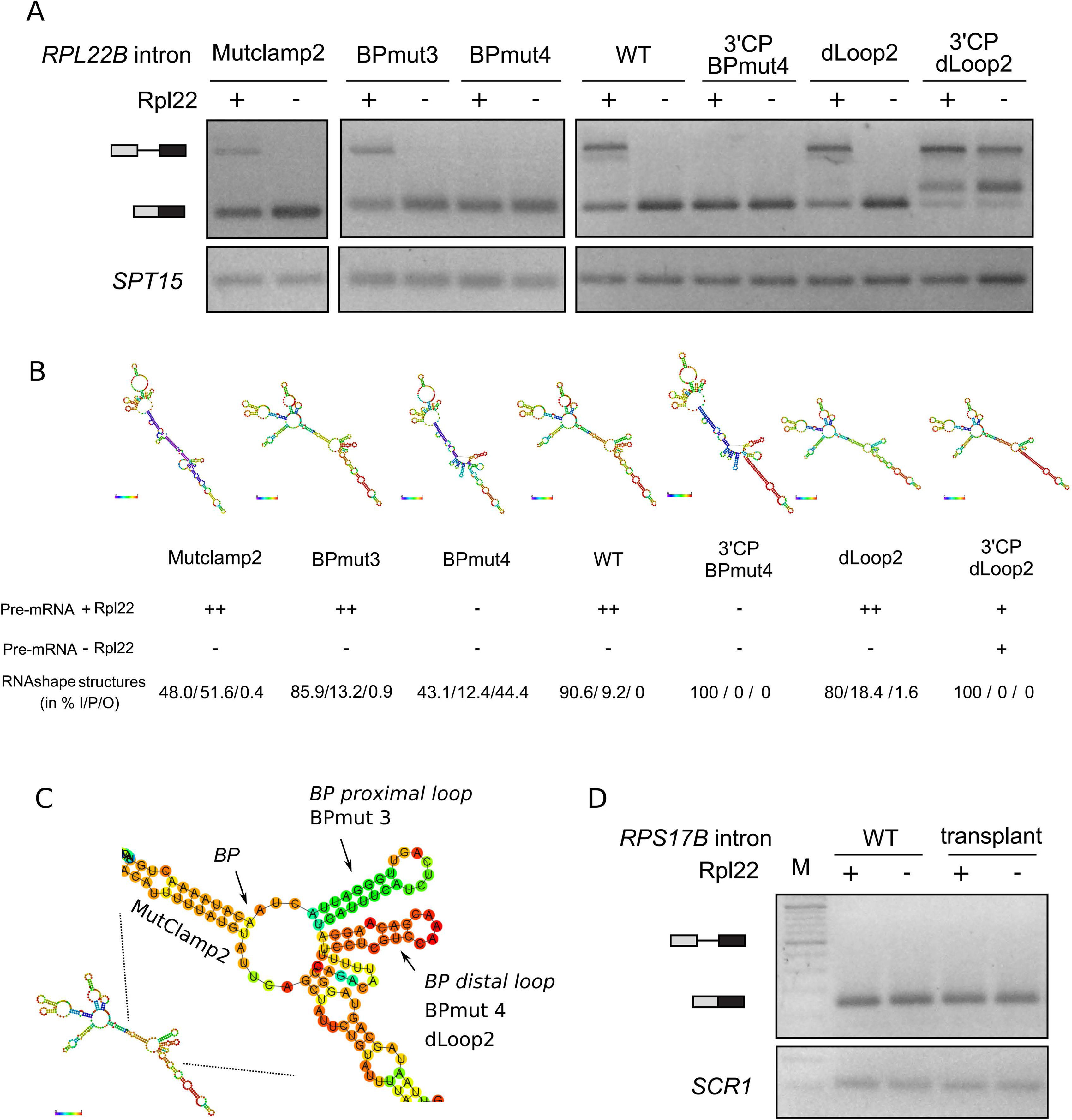
The main stem loop of the inhibitory structure of *RPL22B*i was necessary but not sufficient for splicing inhibition. The experiment was done and the figure was formatted as in Fig 2. **(A) The inhibitory mechanism required that specific structural details of the I structure remain intact.** In the mutants analyzed in this set, the main stem loop is predicted to form. Melting of Loop2 of the I structure (BPmut4; see panel (C)) renders the structure splicing permissive, while ablating the loop (dLoop2) retains the Rpl22-dependent inhibitory potential. Although the stability of the main stem loop should be lowered in Mutclamp2, it is apparently still sufficient to keep the I structure Rpl22 responsive. **(B) Summary of the reporter experiments in (A) and types of stem loop arrangement predicted by RNAshapes.** pre-mRNA accumulation was approximated on a scale of ‘+++’ to ‘-’. The secondary structures were predicted using RNAshapes [54]. The structures were sorted according to the stem loop arrangement involving the conserved region and its complementary elements (see Fig 1), and the number of structure types classified as I, P, or O (other) was expressed in %. See text and Figs. S1-S3 for additional information. **(C) Detail of the predicted secondary structure of *RPL22B*i in the I conformation.** The regions referred to in the text are indicated (in italics) together with the mutants (in plain text). **(D) The main stem loop was not sufficient to convey splicing inhibition in an unrelated intron.** *The RPS17B* intron without or with the main stem loop inserted was tested as in (A). M indicates the m.w. marker lane.

The effects that we observed in the mutants described in Fig 4 cannot be explained solely by the dichotomy of the alternative 5’COMP/CONS versus 3’COMP/CONS base pairing. According to RNAshapes, 3’CP+BPmut4, 3’CP+dLoop2, dLoop2 and Mutclamp2 return >80% I structures, yet the splicing results covered the whole range - from ‘permissive’ to ‘regulated’ to ‘inhibitory’. According to Sfold, BPMut4 and especially 3’CP+BPMut4 maintain the characteristics of an ensemble ‘I’ (S6 Fig), but the efficiency of splicing inhibition was diminished. In the structures of BPMut4 predicted by RNAshapes, we observed distinct arrangements between the main stem loop and the portion containing 5’ss of the secondary structure diagram (Figs. 4B and S4). We hypothesize that while the main stem loop is part of the inhibitory mechanism, the efficacy with which the RNA structure interferes with the spliceosome depends on the details of its 3D contacts. In comparison, partial destabilization of the main stem loop had only a limited effect on splicing inhibition.

Gabunilas and Chanfreau inserted part of the *RPL22B* intron, spanning nucleotides 153 through 239, into the modified intron of *RPS21A*, but did not see an inhibitory effect on splicing after overexpression of Rpl22A [11]. According to our predictions (see Fig 5B), the intron part which the authors used did not contain the main stem loop in its whole. Therefore, we transplanted the full extent of the stem loop (nt 153 to nt 246) into the intron of *RPS17B*. Like *RPL22B*, *RPS17B* is a ribosomal protein gene with a suboptimal 5’ss (GTACGT) and similar intron length (314 nt). The stem loop was inserted at the same distance from the BP as it is located in *RPL22B* (220 nt in the intron), and the construct contained 60 nt of exon2. Stem loop formation in this new context was predicted by RNAfold (S4 Fig). The experiment did not show an inhibitory effect of this stem loop on *RPS17B* splicing (Fig 4D). Although this result does not exclude the possibility that the stem loop might inhibit splicing in a different setting, it supports the conclusion that additional requirements must be met for the inhibition to occur.

**Fig 5.**
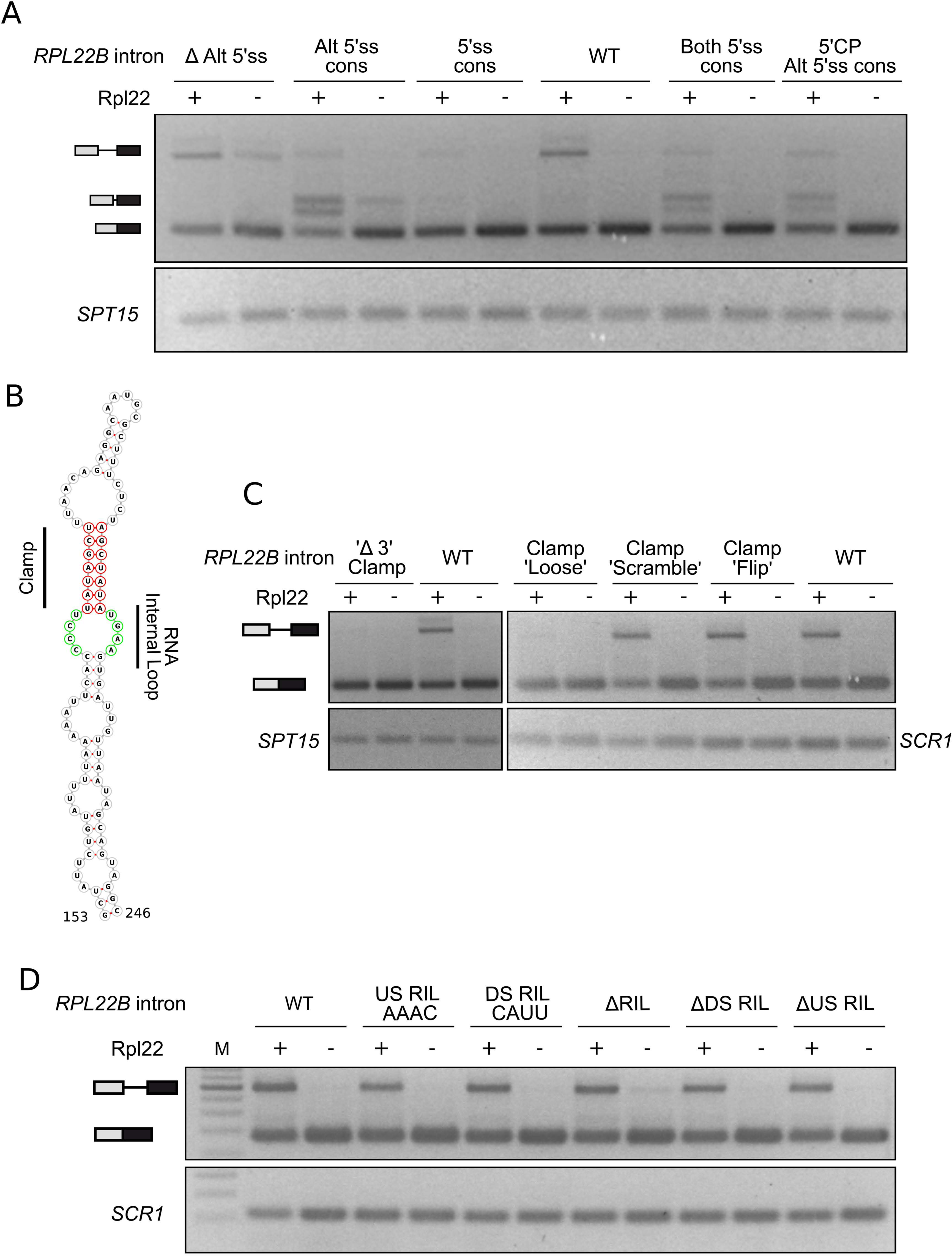
The splicing of the *RPL22B* intron was affected by the pre-mRNA context. The experiment was carried out as described in Fig 2. **(A) Both main and alternative 5’ss contributed to the *RPL22B*i splicing inhibitory mechanism.** The removal of alternative 5’ss increased the level of unspliced RNA in the absence of Rpl22 compared to WT (ΔALT5’ss). Mutating the 5’ss sequences to the yeast consensus decreased the ability of Rpl22 to inhibit splicing and increased the use of alternative 5’ss. The alternative splicing products migrated above the size of the major band and the PCR products of the corresponding length were sequenced to confirm this assumption. **(B) Predicted secondary structure of the main stem loop of *RPL22B*i.** The structure was modeled by RNAfold [52], visualized using Forna [58] and annotated according to [11]. ‘Clamp’ and ‘RNA internal loop’ (RIL) are labeled red and green, respectively. In [11], ‘Clamp’ is referred to as ‘Lower distal stem’. **(C) Structure, but not the sequence of the ‘Clamp’ was important for splicing regulation.** Deletion of the 3’strand of the ‘Clamp’ or weakening of its complementarity rendered intron splicing permissive regardless of Rpl22. On the contrary, maintaining complementarity while scrambling the sequence or flipping the stem arms retained the WT behavior. **(D) Nucleotides within the ‘RNA internal loop’ were dispensable for the regulation of RPL22Bi splicing.** In contradiction to previous findings [11], nucleotides within the ‘RNA internal loop’ (RIL) are dispensable for the regulation of RPL22Bi splicing. CCCU to AAAC mutation in the 5’arm of the loop (‘5’RIL AAAC’), the UGAA to CAUU mutation in its 3’arm (‘3’RIL CAUU’), or the deletion of the loop or its arms did not lose regulation after the overexpression of the Rpl22 protein. ΔRIL indicates the deletion of both 5’arm CCCU and 3’arm UGAA nucleotides; Δ5’RIL and Δ3’RIL indicate the per partes deletions of CCCU and UGAA, respectively.

### *RPL22B* intron splicing is affected by the pre-mRNA context

The 5’ss of *RPL22B*i is non-canonical (GTACGT) and is accompanied by an ‘alternative’ 5’ss (ALT5’ss) 65 bp downstream. The latter site does not produce a translatable ORF and genomic data measured under standard cultivation conditions showed that it was being used only marginally [12,59–62]. We noticed the accumulation of alternatively spliced products in mutants in which the usage of the main 5’ss was inhibited (see Fig 2A). The identification of these molecules was done by sequencing the PCR product of the corresponding length.

We first asked whether the inhibitory mechanism would be capable of overcoming a stronger signal, that is, the canonical 5’ss signal. The mutant with the canonical 5’ss sequence (GTATGT; 5’ss-CONS) showed weakened but still discernible inhibition by Rpl22A (Fig 5A). Next, we mutated ALT5’ss to consensus, which lowered the Rpl22A-induced accumulation of pre-mRNA compared to WT while increasing the accumulation of alternatively spliced product(s). With increased inhibition (i.e. +Rpl22A), the band intensity of the main spliced RNA decreased, while that of the alternative product increased. The effect of consensus ALT5’ss prevailed even in the 5’CP+ALT5’ss-CONS mutant, where the 5’COMP/CONS base pairing was stabilized (and inhibition lessened; see Fig 3). We still observed alternative splice products that were correlated with the expression of Rpl22A. We reasoned that if the inhibition mechanism did not involve the 5’ part of the intron, the ratios (+Rpl22A)/(-Rpl22A) for the spliced products would be similar for both the main and the alternative 5’ss used. Their anticorrelation suggested that both parts of the intron contributed to the inhibitory mechanism.

When we deleted ALT5’ss (deletion of nt 65-70), we observed enhanced inhibition in the absence of Rpl22A (Fig 5A), which further supported the role of the 5’ss recognition in the mechanism. The result showing that the deletion of ALT5’s negatively affects the splicing efficiency of *RPL22B*i was already obtained by the Chanfreau group (Fig 5B in [59]; S2C Fig in [11]). We concluded that the whole intron is important for the execution of the inhibition, either because of its 3D structure or because its parts differentially affect the assembly of the spliceosome.

The formation of the main stem loop, which participates in Rpl22A binding, is a prerequisite for inhibition of splicing. Both the 3’COMP/CONS stem and the shorter clamp stem contribute to the formation of the structure. The removal of the 3’ arm of the clamp stem (Δ3’Clamp; deletion of nt 214-220; see Fig 5B) disrupted inhibition (see Figs. 3 and 5C). To confirm the structural role of the clamp we mutated the stem to decrease its stability (Clamp’Loose’; mutation of nt 186GC to CG and 218AT to TA), scrambled the base pairs while maintaining complementarity (Clamp’Scramble’) and flipped the sequences between the 3’ and 5’ arms of the stem (Clamp’Flip’). Although the Clamp’Loose’ mutant lost its ability to inhibit splicing, the latter two mutants showed WT behavior (see also [11]).

The reporters used by [11] lacked the alternative 5’ splice site as well as the surrounding exons (see S8 Fig for comparison). To address whether these manipulations could have affected the outcome of the assay, we recreated some of their mutants of the ‘RNA internal loop’ (RIL; see Fig 5B) using our approach. We constructed the the CCCU to AAAC mutation in the 5’arm of the loop (‘5’RIL AAAC’), the UGAA to CAUU mutation in its 3’arm (‘3’RIL CAUU’), or deletion of the loop (ΔRIL) or its arms (Δ5’RIL and Δ3’RIL), all of which were in the context of the entire intron. In contrast, the constructs tested by Gabunilas and Chanfreau [11] lacked the apical portion of the stem between nt 191 and 211, which obviously made them fold differently from the full-length stem, irrespective of the deletion of the internal loop. In our hands, none of the above mutations disrupted the negative regulation of splicing by Rpl22A because all reporters reacted to Rpl22A overexpression by increasing the amount of unspliced pre-mRNA to levels obtained in WT (Fig 5D). In the uninhibited state, they were also spliced with the same efficiency as the WT intron; a possible exception was the ΔRIL mutant, which showed slightly less efficient splicing. None of the RIL-targeting mutations impaired the ability of this part of the intron to interact with Rpl22 in Y3H (S7 Fig). This further supports our conclusion that mutations in ‘RNA internal loop’ nucleotides, including its DS arm GUAA (nt 178CCCU181 and 221UGAA224), do not alter Rpl22A-mediated inhibition.

### Splicing inhibition occurs during cotranscriptional spliceosome assembly

Our previous analyses of *RPL22* splicing identified varying degrees of pre-mRNA accumulation depending on the paralogue and the level of Rpl22, but never the lariat-exon2 intermediate, suggesting that inhibition occurs before the first splicing step [12]. Notably also, the splicing sequences were not part of the Rpl22 binding stem loop. We wanted to confirm that the inhibitory mechanism does not involve blocking splice site access but rather targets later stages, that is, the assembly of spliceosomes.

We analyzed the cotranscriptional recruitment of the components of the yeast commitment complex 1 Prp42 (U1 snRNP), Msl5 and Mud2 (BP recognition) to the *RPL22B* locus using ChIP-qPCR analysis as previously described [63]. Cells expressed HA-tagged versions of splicing factors and primers (P to E; Fig 6) were used to amplify DNA fragments along the *RPL22B* locus after chromatin anti-HA immunoprecipitation. We took advantage of the fact that the strain *rpl22a*Δi produces high levels of Rpl22A and compared this strain with the strain *rpl22a*Δ (=no Rpl22A) and the strain ‘WT’. Under the high concentration of Rpl22A (*rpl22a*Δi), splicing of *RPL22B*i is inhibited [12]. We were able to obtain WT recruitment profiles of the tagged factors for both *rpl22a*Δi and *rpl22a*Δ strains (Fig 6). This implies that the regulatory mechanism did not block the first stage of recognition of 5’ss and BP. We were unable to analyze in detail the binding of subsequent spliceosome assembly components due to the short length of exon 2 in *RPL22B*.

**Fig 6.**
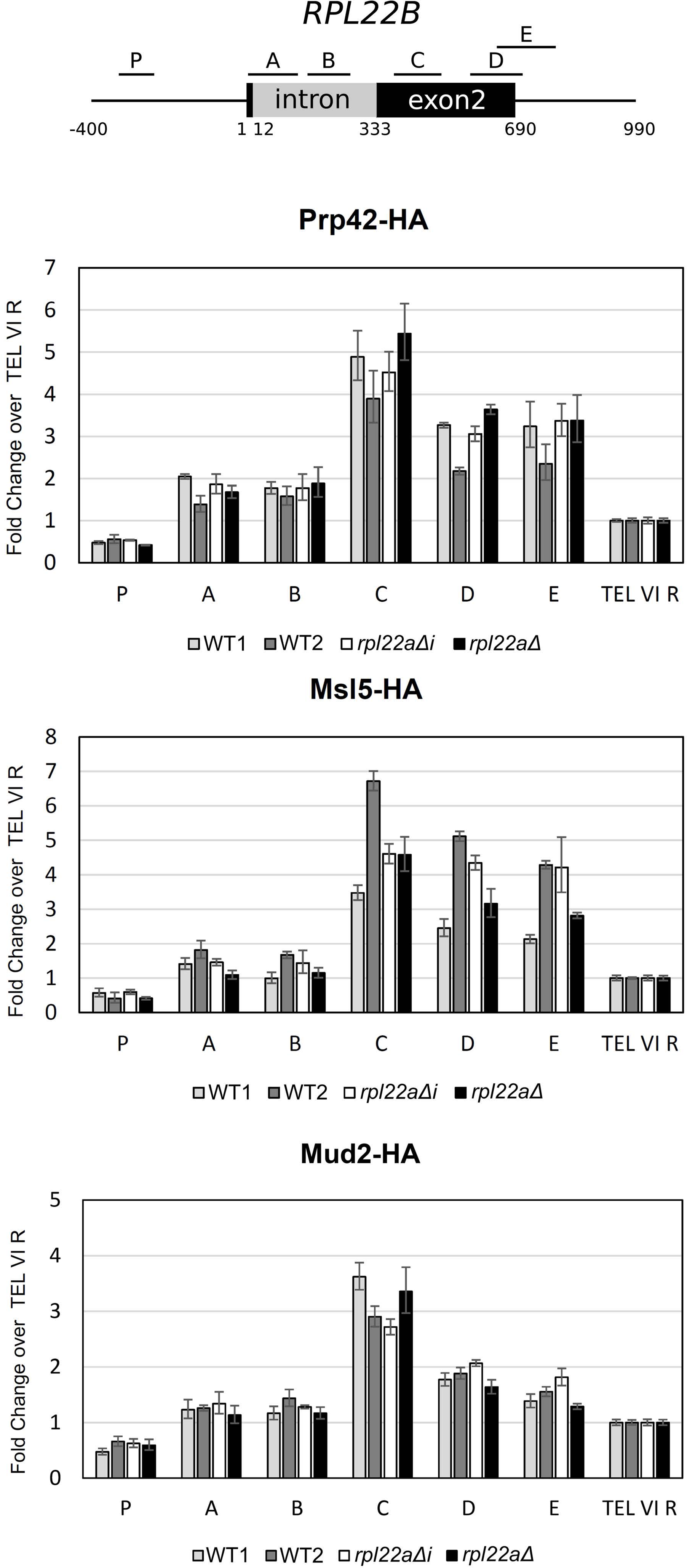
Splicing inhibition did not preclude cotranscriptional recruitment of U1 snRNP and BBP to the *RPL22B* locus. The recruitment of spliceosomal components to the *RPL22B* locus was analyzed by ChIP-qPCR analysis. We used cells expressing HA-tagged Prp42, Msl5, and Mud2 to monitor the cotranscriptionality of the formation of commitment complex 1. The signals were normalized to input values and expressed as the fold enrichment relative to the signal obtained from the telomeric region (TEL VI R). A representative graph of three independent experiments is shown; error bars represent the S.D. of three technical replicates. To compare the effect of low and high concentrations of Rpl22, we used *rpl22a*Δ and *rpl22a*Δi strains, respectively. Positions of the PCR amplicons P to E are indicated.

### The Rpl22 homologs complement the ability of Rpl22A to inhibit splicing in *S. cerevisiae*

The demonstration that *Kluyveromyces lactis RPL22* (*KlRPL22*) can be regulated by changes in the concentration of Rpl22A/B or KlRpl22 proteins [12] led us to ask about the ability of other homologues of Rpl22 to complement the extra ribosomal function of Rpl22A in *S. cerevisiae*. Rpl22 binding is dependent on the presence of lysines in the RNA binding interface of the protein, because mutants in which these lysines were substituted with glutamates were not binding in Y3H and were unable to inhibit *RPL22B* splicing *in vivo* [12]. The conservation of the residues that contribute to charge attraction can be shown in the alignment of Rpl22A with other Rpl22 homologs that we used for our complementation analysis (S8 Fig). We mapped the amino acid changes among the seven selected homologs on the surface of the Rpl22 structure. As expected, the part of the protein that faces the rRNA in the ribosome [64] was the most conserved (Fig 7A).

**Fig 7.**
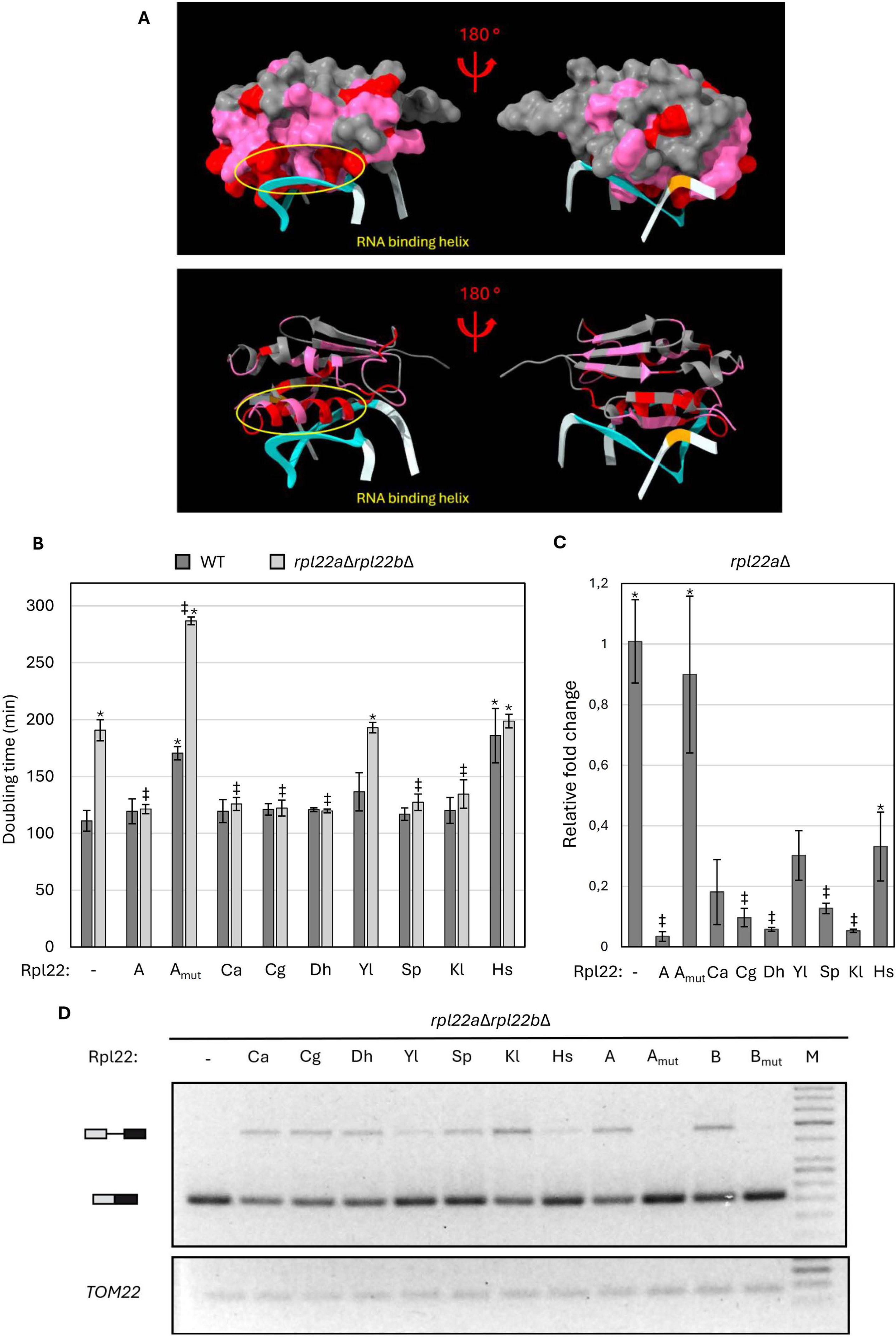
Rpl22 homologs complemented the extra ribosomal function of Rpl22A in *S. cerevisiae*. **(A) Projection of invariant amino acids on the surface of Rpl22A of *S. cerevisiae.*** The structure of Rpl22 and 25S rRNA was extracted from the ribosome structure (PDB: 4V88) [64] using the ChimeraX program (https://www.rbvi.ucsf.edu/chimerax) [65]. Helices 57 and 59 of 25S rRNA are highlighted in blue and orange, respectively. The pooled amino acid differences between Rpl22A and the Rpl22 homologs tested in (B) to (D) are projected onto the surface of the molecule. The colors indicate the identity to Rpl22A (red), conservative substitutions (pink; see ‘:’ and ‘. in the alignment in S8 Fig), and nonconserved positions (gray). The projection delineates parts that are likely involved in the intron-regulatory mechanism of *RPL22*. **(B) The Rpl22 proteins from related yeast species, *S. pombe* and *H. sapiens* were tested for their ability to support WT growth of the *rpl22a*Δ *rpl22b*Δ strain.** We expressed in *rpl22a*Δ *rpl22b*Δ and WT *S. cerevisiae* cells different *RPL22* genes as intronless versions under the control of the *ADH1* promoter (plasmid pVTU260) and evaluated whether the production of different Rpl22 proteins would rescue the growth phenotype. Transformation with the empty pVTU260 served as a control (-). The Rpl22 proteins tested included Rpl22A (A), the binding-dead mutant of Rpl22 (A_mut_) and the homologs from *Candida albicans* (Ca), *Candida glabrata*(Cg), *Debaryomyces hanseni* (Dh), *Yarrowia lipolytica* (Yl), *Schizosaccharomyces pombe* (Sp), *Kluyveromyces lactis* (Kl) and *Homo sapiens* (Hs). The y-axis represents the generation time of individual strains. The plotted values are the means and standard deviations of three biological replicates. The results of the Student’s t-test with Bonferroni correction for multiple tests are indicated by (*) or (‡) for the differences between a strain and WT or the double deletant *pl22a*Δ *rpl22b*Δ, respectively (both (*) and (‡) indicate *P*≤0.001). **(C) Endogenous *RPL22B splicing* can be inhibited by Rpl22 homologs in the *rpl22a*Δ strain.** *The RPL22* homologs were expressed in the *rpl22a*Δ strain as in (B). After the preparation of cDNA from exponentially growing cells, the level of endogenous *RPL22B* mRNA was measured by qPCR using primers specific for the *RPL22B* locus. The signal was normalized to *SPT15* and to the *rpl22a*Δ strain transformed by an empty vector (-). Splicing inhibition was caused by the expression of *S. cerevisiae* Rpl22A (A) but not its binding-dead mutant (A_mut_). The Rpl22 homologs were labeled as in (B). Statistical significance of the difference between the *rpl22a*Δ strain harboring an empty vector (-) and a Rpl22 expressor is indicated as (‡); (*) is used for the difference between (A) and other strains. Both symbols indicate *P*≤ 0.005 based on a t-test with Bonferroni correction for multiple comparisons. **(D) All tested Rpl22 proteins can inhibit splicing of the *RPL22B* reporter in vivo.** The *rpl22a*Δ *rpl22b*Δ cells expressing the indicated homologues of Rpl22 (see (B)) were transformed with a plasmid carrying the *reporter RPL22B-CUP1.* The reporter splicing efficiency was measured and the figure was formatted as in Fig 2. *TOM22* was used as a loading control. The figure represents the result of one of three independent experiments.

We used the *rpl22a*Δ *rpl22b*Δ strain of *S. cerevisiae* and first tested the ability of the Rpl22 homologues to complement the ribosomal function of endogenous Rpl22A/B. The deletion of both *RPL22* genes *was* shown to result in 1.7 times slower growth compared to WT *S. cerevisiae* [66]. Upon expression of Rpl22A, the doubling time was reduced to WT level (Fig 7B). The expression of Rpl22 homologs from *Candida albicans*, *Candida glabrata*, *Debaryomyces hanseni*, *Kluyveromyces lactis*, or *Schizosaccharomyces pombe* had a similar effect. The exceptions were the proteins of *Yarrowia lipolytica* and *Homo sapiens*. While these proteins were expressed at detectable levels in the *rpl22a*Δ *rpl22bΔ* strain (S10 Fig), they were unable to reduce the doubling times. The expression of human Rpl22 increased the doubling time of WT, similarly to the effect of the binding-dead mutant of Rpl22A (A_mut_). We assume that the toxicity resulted from interference with normal ribosome function.

To see the effects of the homologs on *RPL22* splicing, we measured the levels of endogenous *RPL22B* mRNA in a *rpl22a*Δ strain by qPCR. All tested proteins, except for Rpl22mut, decreased the levels of *RPL22B* mRNA (Fig 7C). The effects of the homologs of *C. glabrata*, *D. hanseni*, *K. lactis*, and *S. pombe* inhibited splicing comparably to Rpl22A of *S. cerevisiae*. The effects of *Y. lipolytica* and human homologs (< 40% of the empty vector strain) could have been due to lower protein levels (S10 Fig) and/or toxicity. In addition, we also analyzed inhibition of *RPL22B* splicing using our *RPL22B-CUP1* reporter system. The experimental setup was described in Figs. 2-4. The reporter showed no trace of unspliced product when the binding-dead mutant was expressed, in contrast to the effects of *S. cerevisiae* Rpl22A/B, which led to pre-mRNA accumulation. Among the homologs, *Y. lipolytica* and human products showed the weakest effects, but all the proteins tested inhibited splicing to some extent (Fig 7C).

The map of the amino acid differences among the Rpl22 homologs shows that while the rRNA interacting helix contains only conservative substitutions, the opposite side of the protein harbors non-conservative changes (Fig 7A). Because both the ribosomal and extra ribosomal functions of Rpl22A were complemented by the various homologs, it is likely that these functions use the same interface on the protein.

## Discussion

### The inhibitory mechanism requires the intron as a whole

We used mutagenesis of the *RPL22* introns to test whether their splicing efficiency depends on the formation of a particular RNA structure (as predicted by RNA fold and other algorithms). Our experimental setup allowed us to differentiate mutations that were splicing inhibitory regardless of the presence or absence of the Rpl22 protein (see Fig 2) from those that were permissive in its presence, rendering the intron unresponsive to Rpl22-mediated inhibition (see Fig 3). Furthermore, some mutations showed graded effects and some affected the relative use of ALT5’ss. We grouped mutants into three categories according to the predicted structures they contained, those with (i) stabilized inhibitory conformations (I), (ii) stabilized permissive conformations (P) and (iii) ‘disabled’ (= splicing permissive) I conformations. In contrast to the previous study [11], we retained ALT5’ss and included intron proximal sequences in our constructs. This was because we found that the deletion of *RPL22B*i ALT5’ss impaired the splicing efficiency at the major site (Fig 5A). The *in vivo* splicing of *RPL22B* uses alternative 5’ and 3’ splice sites, albeit at low frequencies [12,59–62]. Although ALT5’ss does not lead to a functional product, it may be part of the spliceosome assembly process, for example, as an adjunct assembly landing platform [37,67].

Among the three categories of mutations, the ‘disabled’ I conformations were the most interesting, as they apparently targeted the inhibitory mechanism outside of the main stem loop, which was nevertheless still predicted to form (BPmut4 and 3’CP+BPmut4; see Fig 4). The ensemble of secondary structures of BPmut4 was predicted to be similar to WT (I>P), but the splicing *in vivo* was not blocked. We concluded that the predicted formation of the stem loop itself was not sufficient for splicing inhibition. We cannot exclude that in this mutant the binding of Rpl22 *in vivo* was not affected even though the motif that bound the protein in Y3H was intact. There may be a part of the RNA structure other than the main stem loop that contacts Rpl22. However, the destabilization of Rpl22 binding in other mutants (= nonbinding Rpl22 in Y3H) did not automatically result in splicing permittivity. This is why we assumed that the defect in BPmut4 was not related to the absence of Rpl22 binding. We speculate that the defect in inhibition of splicing in BPMut4 was due to (i) changes in 3D contacts within the intron structure or (ii) changes in the energy barrier that separates I and P conformations. We noticed that the predicted conformations of BPmut4 differed from those of WT in details that included the 5’COMP base pairing and the nucleotides between 5’COMP and 3’COMP. Structure predictions postulate two stem loops 5’ proximal to BP (see Fig 4C). The melting of the more proximal of the two (in BPmut3) did not affect the inhibition outcome, while melting of the more distal one did (in BPmut4). Moreover, ablating this latter loop kept the inhibitory effect intact. If the I/P ratio was under kinetic control, BPmut4 can work by lowering the energy barrier that separates the I and P conformations and thus allowing more P structures to form cotranscriptionally (and splice). We concluded that blocking BP access is not the way the mechanism operates and that it relies on features throughout the intron, including the 5’ splice sites.

Base pairing of conserved elements across introns was previously described as a means of enhancing splicing in *RPS17B* and several other RPGs [37,68–70]. Rogic and co-workers tried to demonstrate that a ‘structural distance’ between 5ss and BP, defined as the shortest walk on a 2D map of secondary structure prediction, was correlated with splicing efficiency. Considering the ‘structural distances’ between 5ss and BP in the conformations of *RPL22B*i, we found that they cannot explain the inhibition observed in *RPL22*. In the I conformation, ca 100 to 180 nucleotides are ‘removed’ by stem loops (depending on the stability of the stem), whereby the 5’ss BP distance is thus ca 40 to 50 nucleotides (measuring a shortest path on a graph of a secondary structure). Although the P conformation orients the 5’ss and BP in closer proximity, neither of the two conformations should pose a challenge of distance. Spliceosome assembly proceeds co-transcriptionally, with the U1 snRNP complex being anchored by the cap binding complex and the PolII holocomplex [71–74]. The BP recognition complex is also in contact with Prp40, Prp39 and Nam8 of U1 snRNP and genetically interacts with the *TGS1* cap binding complex enzyme [75–78]. Without some obstacle effect that prevents alignment of the 5’ss and BP binding complexes, the assembly of the spliceosome would not be blocked.

### *Inhibition of RPL22B*i does not block recognition of 5’ss and BP

The cotranscriptional recruitment profiles of the U1 snRNP component Prp42 (see Fig 6) and the heterodimer BBP/Mud2 were not affected by splicing inhibition, suggesting that the steps further downstream in the spliceosome assembly pathway were responsible for the blockage. We presume that the critical steps lie between the formation of the prespliceosome A and the maturation to the spliceosomes pre-B and B [76,79,80]. Gabunilas and Chanfreau, who also obtained ChIP data for Prp42 recruitment at the *RPL22B* locus, concluded that they found partial inhibition of Prp42 binding, which may not fully explain the observed inhibition of splicing. Indeed, their ChIP results (Fig 4G in [11]) showed only moderate signal reduction. An interesting comparison can be made with inhibition of *RPL30* splicing by the Rpl30 protein [81,82]. Rpl30 binds a kink-turn structure that encompasses the 5’ss of its own pre-mRNA and thus inhibits splicing. Inhibition did not block U1 snRNP and BBP/Mud2 recruitment, but affected U2 snRNP binding, likely also blocking subsequent rearrangement steps.

Although the inhibitory mechanism did not block the first step of recognition of 5’ss, the result of inhibition was dependent on the presence of ALT5’ss and the strength of both the main and alternative splice sites. A canonical main 5’ss made the inhibition less efficient (Fig 5A). The non-canonical sequence 5’GUAC**GU**(U) contains U at the 7th position, which should weaken the interaction with U6 snRNA (through the 5’caau**AC**a motif of U6). In yeast splice sites, C at the 4th position was found to be correlated with A at the 7th position [83]. In ALT5’ss GUU**UGU**(C), the base pairing with U6 would be canonical, but the interaction with U1 would be weaker. Perhaps this is part of the reason why the main and alternative sites behave distinctly with respect to Rpl22-induced inhibition.

### Rpl22 homologs complement the ability of Rpl22A to inhibit splicing in *S. cerevisiae*

The Rpl22 proteins of other yeasts, as well as human RPL22, complemented the inhibitory function of Rpl22A (see Fig 7). The Rpl22 homologs were constrained in evolution to bind to highly conserved rRNA and function in the ribosome context [84]. As we did not find any example of an Rpl22 homolog that did not inhibit the splicing of *RPL22B* at least partially, it is highly probable that the extra ribosomal function of Rpl22 uses the protein’s rRNA binding site. At the same time, given that only 26 out of 121 amino acids of Rpl22A are identical among the seven homologs, the interaction seems to be flexible. The imperfect stem of the main stem loop of *RPL22* intron contains a motif that is identical in both paralogs as well as all *RPL22* genes of the species of the Saccharomycotina group (Fig 1A). The proteins accommodate both the 25S rRNA loops [64] and the intronic RNA without any apparent similarity between the two. In higher eukaryotes, but not in yeast, some of the RNAs whose expression is regulated by Rpl22 contain a consensus hairpin that fits the Rpl22-interacting part of 25S rRNA [22].

### *The RPL22* intron behaves as an RNA switch

According to cross-linking experiments and structural information, intron - spliceosome contacts are not limited to the splice sites themselves but include additional sequences around them [85,86]. A striking physical association was documented between an intronic stem placed between 5’ss and BP and a spliceosome in the *ACT1* intron [77,86]. The stem loop was placed across positively charged residues of Prp39, Prp42, and the C-terminal domain of U1C as part of the E complex. Mutations destabilizing the stem loop led to pre-mRNA accumulation of the reporter gene, suggesting that interactions via this stem loop aided splicing efficiency [77].

The spliceosomal RNP complex goes through a series of conformational rearrangements before it can catalyze the first transesterification step. Intronic structures could affect these processes if they form anywhere within the footprint of the assembling spliceosome [87]. Kumar and coauthors performed a structure-guided prediction analysis of the effects of mutations on alternative splicing of the exon 10 - intron 10 junction of the MAPT gene. They obtained the best prediction accuracy when considering a footprint of 43 nucleotides for the B^act^ complex. The authors argued that the RNA structure “functions as a rheostat” in this range of nucleotides, which must be unfolded to participate in the spliceosome - pre-mRNA interface.

According to the prediction of Boltzmann weighted structural ensembles for *RPL22*i, the tendency to form I conformations is higher in *RPL22B*i than in *RPL22A*i (S6 Fig). In fact, endogenous *RPL22B* was more stringently regulated than *RPL22A* - after Rpl22 overexpression, *RPL22B* mRNA production dropped to ∼10% [12,88]. We hypothesize that a nascent pre-mRNA structure of *RPL22* interacts with the assembling spliceosome and prevents its own splicing. Structures form as the RNA exits the PolII complex [50], suggesting a kinetic control. As splice sites emerge during transcription, they are recognized by U1 snRNP and BBP/Mud2. By the time the 3’ss proximal part is synthesized, the 3’COMP/CONS (= inhibitory) or the 5’COMP/CONS (= permissive) stem loops prevail, depending on the level of free Rpl22. Rpl22 may act here as a factor tuning the I/P ratio through controlling the energy barrier that separates the two conformations (see Figs. 1 and S1). The I/P ratio, in turn, dictates the degree to which splicing is inhibited (and the pre-mRNA degraded). The results of our splicing assays suggest that the inhibition requires the presence of both the main and alternative 5’ss and that it depends on the noncanonical character of the 5’ss. Hypothetically, the presence of the inhibitory RNA structure (I) could lead to aberrant pre-mRNA interactions with snRNAs and / or recruit disassembly factors to spliceosome on an otherwise slow substrate. The spliceosome disassembly process was recently shown to target aberrant B^act^ conformations and to be part of a proofreading mechanism of incorrectly interacting pre-mRNAs [89,90].

The features of the *RPL22* intron invite comparison with the riboswitch mechanism [91]. Binding of small molecules induces a conformational change of their aptamers, which is then transduced as an output via the regulatory platform (Fig 8). An intriguing example of a riboswitch that recognizes a large molecule (tRNA) and represents an ensemble of elongated structures encompassing ∼300 nucleotides, is the T-box riboswitch [92,93]. An example of an RNA switch regulated by protein ligands is the ‘stress responsive RNA switch’ in the 3’untranslated region of mRNA for human vascular endothelial factor A [94]. The switch responds to changing levels of the hnRNP L and GAIT complexes to regulate vascular endothelial factor A expression in response to hypoxia signaling [95,96].

**Fig 8.**
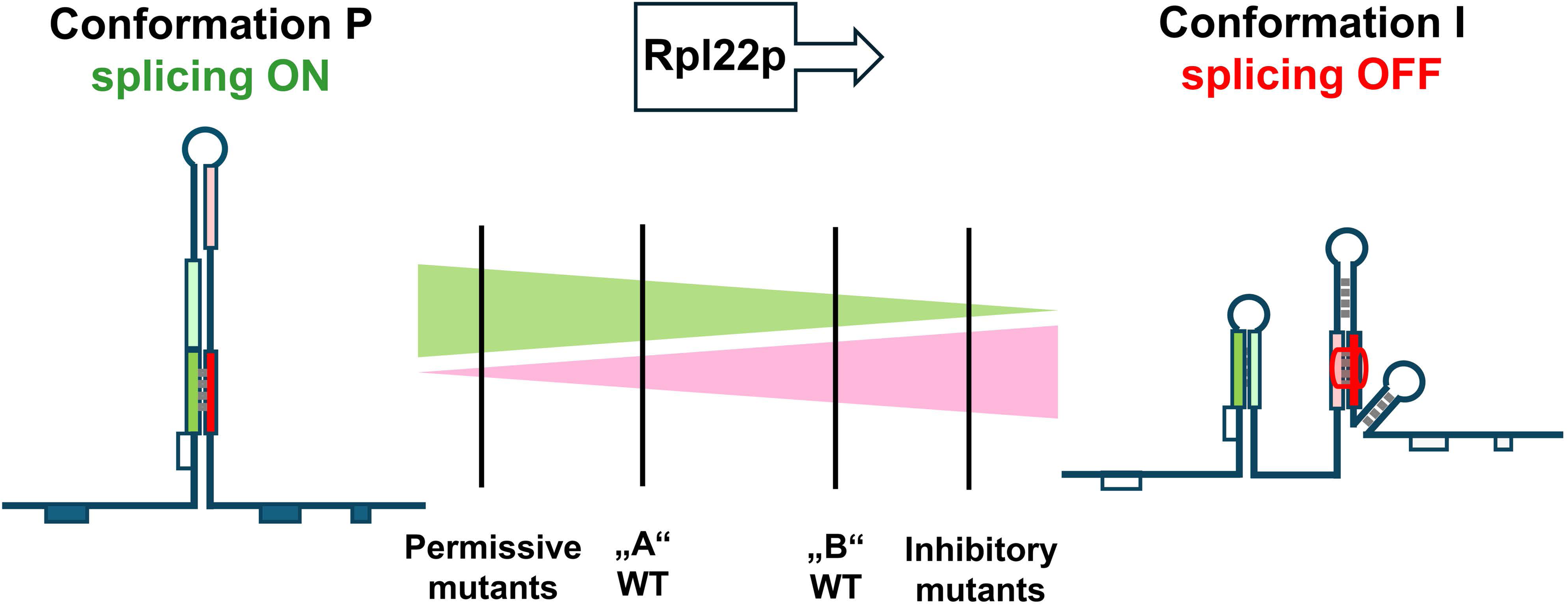
Model of toggling between permissive (P) and inhibitory (I) conformations in *RPLL22* pre-mRNAs. 5’COMP/CONS stem (in P) is mutually exclusive with 3’COMP/CONS stem (in I); drawn in analogy to U2 stem II toggling model [97]. The I conformation inhibits splicing. The effects of mutants and Rpl22p binding are indicated.

Alternate RNA structures may serve as an interface between the inputs, e.g., transcription speed or activity of RNA binding proteins, and the outputs, e.g., alternative splicing or translation. Transcription and transcript processing complexes can act as chaperones, selecting alternate structures from ensembles in response to environmental stimuli [48]. As Saldi pointed out, cotranscriptional RNA folding may be one of the means by which to regulate posttranscriptional splicing [47]. The *RPL22*i - Rpl22 feedback loop represents an intriguing example of an intronic structure, stabilized by the Rpl22 protein, which hinders its own splicing. If confirmed by structural biology (see also [86]), the *RPL22* toolkit of *S. cerevisiae* should help our understanding of how splicing responds to regulatory cues.

## Materials and Methods

### Yeast strains and cultivation methods

Details of the yeast strains used here are provided in Table S1. Yeasts were transformed using the lithium acetate method [98]. The primers and plasmids utilized in this work are listed in Tables S2 and S3. For the splicing assays, yeast cells were inoculated from overnight grown precultures and let to grow for two generations in synthetic medium without uracil and histidine. For the measurement of the doubling times, cells were cultured in medium lacking uracil and tryptophan at 30° C. Optical densities of cell suspensions were recorded spectrophotometrically using Varioskan LUX (λ=600 nm). The analysis of the yeast three-hybrid system was performed essentially as previously described [12].

### Splicing Analysis

RNA isolation and reverse transcription were performed as previously described [12]. Semiquantitative PCR was performed in 25 μl-reactions with 5 μl of 50 times diluted cDNA as template and with primers listed in Table 2 for 25 to 28 cycles. The whole sample was loaded onto 2.5% agarose gel and DNA was stained with ethidium bromide. The photos were taken with a Panasonic DMC-FZ7 camera, processed in the GNU Image Manipulation Program 2.10.6 (https://www.gimp.org) and assembled in Inkscape 0.92.3 (https://www.inkscape.org).

### Plasmid preparation

The intron mutants of *RPL22B*/*A* were synthesized by GeneArt (Thermo Fisher Scientific) and swapped using the restriction sites BamHI and EcoRI with the WT version of *RPL22B* in p423GPD-RPL22B-CUP1. For the three hybrid tests, the mutated versions of *RPL22B* were amplified from GeneArt DNA Strings with primers specific for each mutant (Table S2) and cloned at the SphI site in the vector p3HR2. Plasmids pVTU260 expressing *RPL22* genes from different organisms were prepared as described in [12] using cDNAs as templates for PCR amplification. The cloning outcome was verified by restriction analysis and sequencing.

### ChIP

For ChIP analysis, *rpl22a*Δ and *rpl22a*Δi cells containing a HA-tagged version of a particular spliceosomal subunit (Prp42, Msl5, or Mud2) were cultivated in YPD medium. Precultures were grown overnight and then reinoculated in 50 ml of fresh YPD medium and cultivated for 2-3 generations to reach the mid-log phase. After harvesting, all the steps were performed exactly as described [63]. The DNA fragments were then amplified with primers covering various parts of the *RPL22B* locus using HOT FIREPol® Evagreen® qPCR Supermix on LightCycler 480 II (Roche).

## Acknowledgements

The authors appreciate the excellent technical assistance of Eva Krellerová. We thank Hana Sychrová (Institute of Microbiology of the Czech Academy of Sciences) for kindly providing us the strains used for cloning the *RPL22* genes from *C. albicans* (SC5314), *C. glabrata* (ATCC2001), *D. hanseni* (CBS767) and *Y. lipolytica* (CBS7504). We also thank Martin Převorovský for providing us with the *S. pombe RPL22*. We thank Jiří Libus for constructing the plasmids pVTU260_H. sapiens RPL22 and pACT2_H. sapiens RPL22. We acknowledge the work of Filip Nemčko in the preparation of the strains used for ChIP analyses.

## Abbreviations

3’COMP: 3’complementary element
3’ss: 3’splice site
5’COMP: 5’complementary element
5’ss: 5’splice site
5’UTR: 5’untranslated region
ALT5’ss: alternative 5’splice site
BP: branch point
CONS: conserved sequence element
I2: stem loop region of *RPL22B* intron used in Y3H (nt 165 to 236; Abrhámová et al. 2018)
MFE: minimum free energy
RP: ribosomal protein
RPG: ribosomal protein coding gene
*RPL22B*i: *RPL22B* intron
SD: standard deviation
Y3H: yeast three-hybrid system

## Supporting information

**S1 Fig I ⇋ P interconversion of the intronic RNA would require an extensive change of base pairs.**

**S2 Fig RNAfold predicted structures of introns and their mutants tested in Fig. 2**.

**S3 Fig RNAfold predicted structures of introns and their mutants tested in Fig. 3**.

**S4 Fig RNAfold predicted structures of introns and their mutants tested in Fig. 4**.

**S5 Fig RNAfold predicted structures of introns and their mutants tested in Fig. 5**.

**S6 Fig Base pairing probabilities of Sfold predicted RNA ensembles of *RPL22* introns.**

**S7 Fig Yeast three-hybrid analyses of Rpl22A binding to the I2 fragment of the intronic *RPL22B* RNA.**

**S8 Fig Sequence alignment of *RPL22* homologs tested in this study.**

**S9 Fig The reporter constructs used to analyze RPL22B splicing *in vivo*.**

**S10 Fig Levels of heterologous Rpl22 proteins in the experiments of Fig. 7**.

**S11 Fig Raw images of gels and blots.**

**S1 Table Strains used in this study.**

**S2 Table Primers used in this study.**

**S3 Table Plasmids used in this study.**

**S4 Table List of intronic manipulations tested.**

