## Supplementary material for "Intronic RNA of yeast *RPL22* paralogs acts as an allosteric switch": Supplementary information.pdf

### Supporting information

**Content:**

Supplemental legends and references to supplemental files

Figures S2 to S10

#### Supplemental legends and references to supplemental files

**S1 Fig I ⇋ P interconversion of the intronic RNA would require an extensive change of base pairs.** The I and P conformations of the *RPL22B* intron differ in >20 high probability base pairs as predicted by RNAfold and RNAShapes [1,2]. The animated gif generated using RNA movies tool is shown for illustration [3].

**S2 Fig RNAfold predicted structures of introns and their mutants tested in Fig. 2.** Secondary structures diagrams were generated in RNAfold. The extent of the RNA molecules used for modeling spanned 44 nt in front of the intron and 41 nt after the last nucleotide of the intron. The coloring corresponds to the MFE values indicated under the color bar.

**S3 Fig RNAfold predicted structures of introns and their mutants tested in Fig. 3.** See legend to Fig. S2.

**S4 Fig RNAfold predicted structures of introns and their mutants tested in Fig. 4.** See legend to Fig. S2.

**S5 Fig RNAfold predicted structures of introns and their mutants tested in Fig. 5.** See legend to Fig. S2.

**S6 Fig Base pairing probabilities of Sfold predicted RNA ensembles of *RPL22* introns.** Two dimensional diagrams of base pairing probabilities from a statistical sample of structures were generated by Sfold (*S*rna; [4]) for *RPL22B* WT **(A)**, 3'CW **(B)**, 5'CP **(C)**, BPMut4 **(D)**, 3'CP+BPMut4, and *RPL22A* WT **(D)**. The extent of the sequence used for modeling was the same as in Figs. S2-S5. The position of the apical end of the clamp in *RPL22B* is marked by dashed lines. The densities corresponding to the 3'COMP/CONS (I) and 5'COMP/CONS (P) base pairing are indicated by the red and green arrows, respectively.

**S7 Fig Yeast three-hybrid analyses of Rpl22A binding to the I2 fragment of the intronic *RPL22B* RNA.**

**(A)** The intronic region of *RPL22B* between nt 165 and 236 was tested for its ability to interact with the Rpl22 protein in a yeast three-hybrid assay. Ten-fold serial dilutions of cells were spotted on plates with increasing concentrations of the metabolic retarder 3-aminotriazole (3-AT). '-U', '-L', and '-H' denote the absence of uracil, leucine, and histidine in the medium. Cells with activation domain expressed alone (AD only; right parts of the panels) did not grow in the absence of histidine. Positive (IRE+IRP; [5]) and negative controls (Rpl22-AD with empty plasmid p3HR2) are placed in the last row below the red line. The RNA internal loop mutants supported cell growth to the same extent as the construct containing the WT intron at 1 to 5 mM 3-aminotriazole.

**(B)** Predictions of stem loop formation in the I2 region of the *RPL22B* intron (RNAfold; [1]). The extent of the RNA molecules used for modeling spanned 44 nt in front of the intron and 41 nt after the last nucleotide of the intron. The fragment spanning nucleotides 165 and 236 is shown. For each mutant, two structures are shown: (i) the predicted structure (left; nucleotide color reflects the probability of the particular arrangement) and (ii) the WT template (in grayscale) with manipulated nucleotides highlighted in red (for splicing inhibitory mutants) and green (for splicing permissive manipulations).

**(C)** Yeast three hybrid analysis of MutConsComp1 was done as in (A). Prediction of stem loop formation is shown to the right.

**S8 Fig Sequence alignment of *RPL22* homologs tested in this study.** Rpl22 sequences are shown from top to bottom: *Saccharomyces cerevisiae* Rpl22A (ScRpl22A), *Saccharomyces cerevisiae* Rpl22B (ScRpl22B), *Kluyveromyces lactis* Rpl22 (KlRpl22), *Candida glabrata* Rpl22 (CgRpl22), *Candida albicans* Rpl22 (CaRpl22), *Schizosaccharomyces pombe* Rpl22 (SpRpl22), *Debaryomyces hansenii* Rpl22 (DhRpl22), *Homo sapiens* Rpl22 (HsRpl22) and *Yarrowia lipolytica* Rpl22 (YlRpl22). The segment corresponding to the RNA binding helix is highlighted in yellow. Lysines that were mutated to glutamates in $Rpl22A_{mut}$ and $Rpl22B_{mut}$ are highlighted in red. Asterisks (*) indicate identical amino acids at a given position; colons (:) and periods (.) indicate strong and weak amino acid conservation, respectively. The alignment was generated using Clustal Omega MSA (https://www.ebi.ac.uk/Tools/msa/clustalo/).

**S9 Fig The reporter constructs used to analyze RPL22B splicing *in vivo*.**

**(A)** In this study, we used reporter splicing constructs based on the p423GPD backbone with *RPL22B* intron surrounded by its own 5'UTR, exon 1 and 60 nts of exon 2, which were fused in frame with the *CUP1* reporter. Alternative 5'ss was preserved in all constructs unless indicated. **(B)** In the study of Gabunilas and Chanfreau [6], the reporter plasmid was derived from pUG23. *The RPL22B* intron was inserted in front of the *GFP* reporter without the surrounding sequences. In most of the constructs analyzed, the alternative 5'ss was deleted.

**S10 Fig Levels of heterologous Rpl22 proteins in the experiments of Fig. 7.**

Protein extracts were prepared from exponentially growing cells of the analyzed strains, separated on 10% tris-tricine gels, and transferred to cellulose membranes. Panels show the results of Western blot detection along with the corresponding tris-tricine gels after transfer. Immunodetection of 6xHis-tagged Rpl22 was performed using a primary antibody against the 6xHis tag (6xHis Tag Monoclonal Antibody HIS.H8; Thermo Scientific) and a secondary antibody conjugated to alkaline phosphatase (Goat Anti-Mouse IgG (H+L) AP Conjugate; BIORAD).

**(A) Growth rate complementation experiment in Fig. 7B.** Different *RPL22* genes were expressed as intronless versions under the control of the *ADH1* promoter (plasmid pVTU260) in *rpl22a*Δ *rpl22b*Δ and WT *S. cerevisiae cells* and evaluated. The Rpl22 proteins tested included Rpl22A (A), the binding-dead mutant of Rpl22 ($A_{mut}$) and the homologs from *Candida albicans* (Ca), *Candida glabrata* (Cg), *Debaryomyces hanseni* (Dh), *Yarrowia lipolytica* (Yl), *Schizosaccharomyces pombe* (Sp), *Kluyveromyces lactis* (Kl) and *Homo sapiens* (Hs). The expected molecular weight of the detected protein was approximately 15 kDa (indicated by arrows). Protein extracts from cells carrying the empty pVTU260 vector were used as a negative control (–).

**(B) Splicing analysis experiment in Fig. 7C.** The *RPL22* homologs were expressed in the *rpl22a*Δ strain as in (A).

**(C) qPCR measurement of *RPL22B* mRNA in Fig. 7D.** The *RPL22* homologs were expressed in the *rpl22a*Δ *rpl22b*Δ background as in (A). Negative controls included extracts from cells containing the empty pVTU260 vector and the reporter plasmid p423GPD_*RPL22B-CUP1*.

**S11 Fig Raw images of gels and blots.**

**S1 Table Strains used in this study.**

**S2 Table   Primers used in this study.**

**S3 Table   Plasmids used in this study.**

**S4 Table   List of intronic manipulations tested.**

*RPL22B* intron

S2 Fig

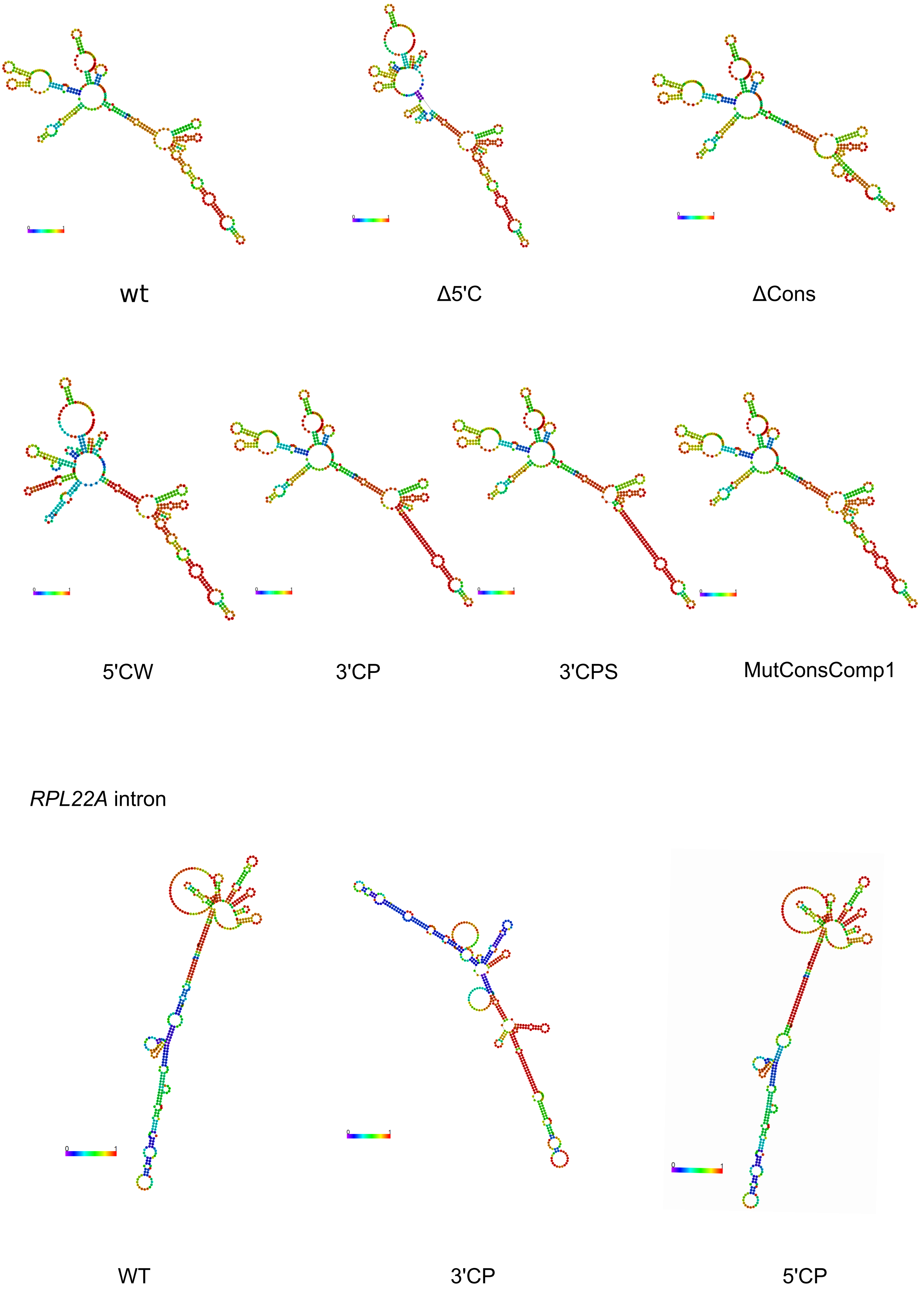

### S3 Fig

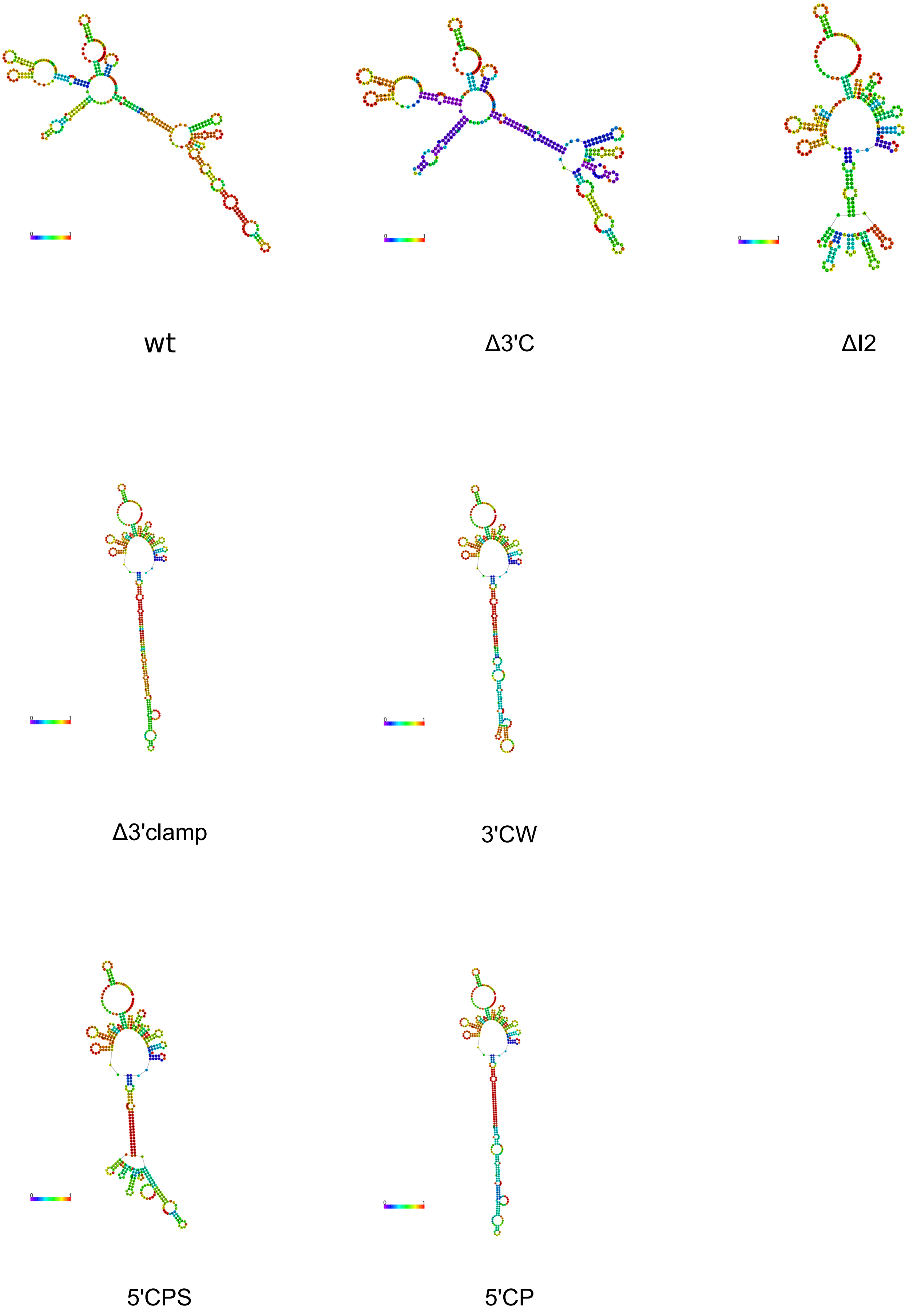

S4 Fig

*RPL22B* intron

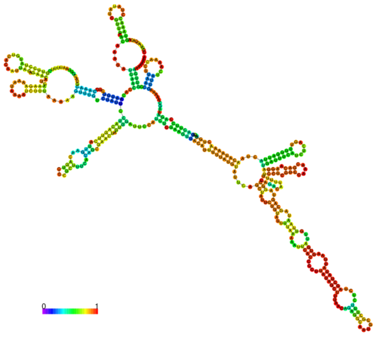

WT

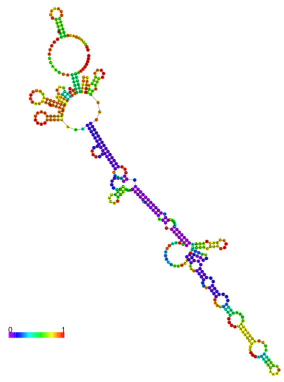

Mutclamp2

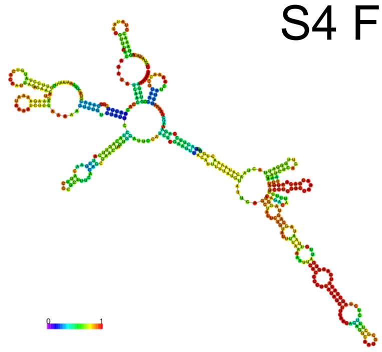

BPmut3

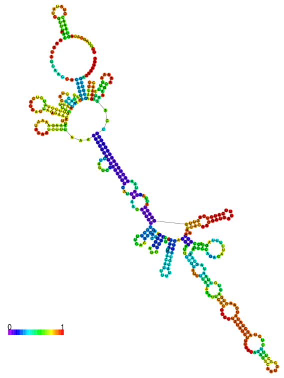

BPmut4

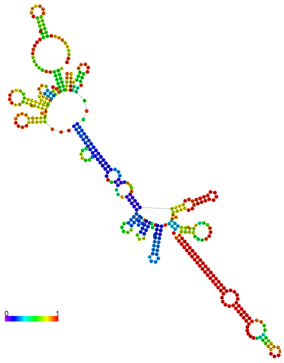

3CP
BPmut4

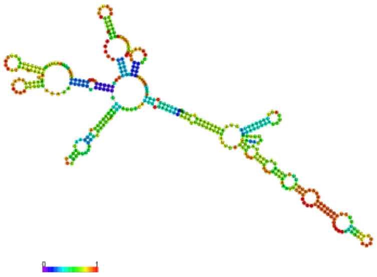

dLoop2

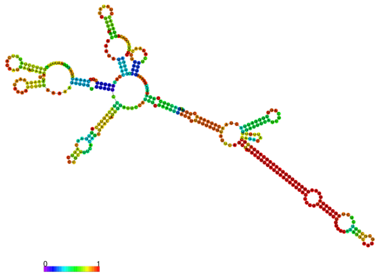

3CP
dLoop2

*RPS17B* intron

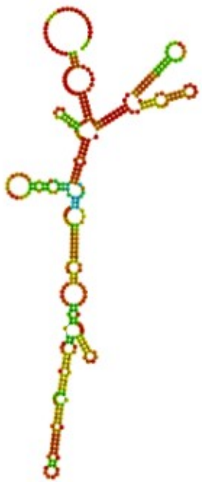

WT

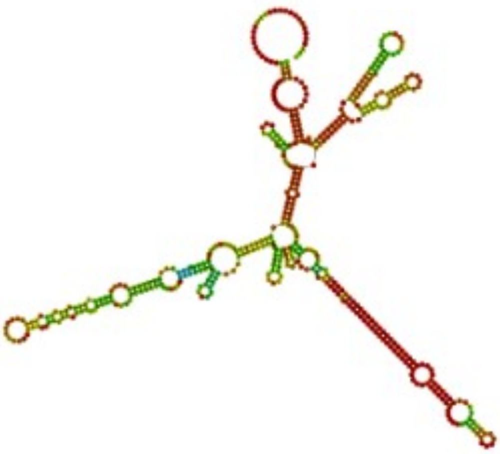

transplant

### S5 Fig

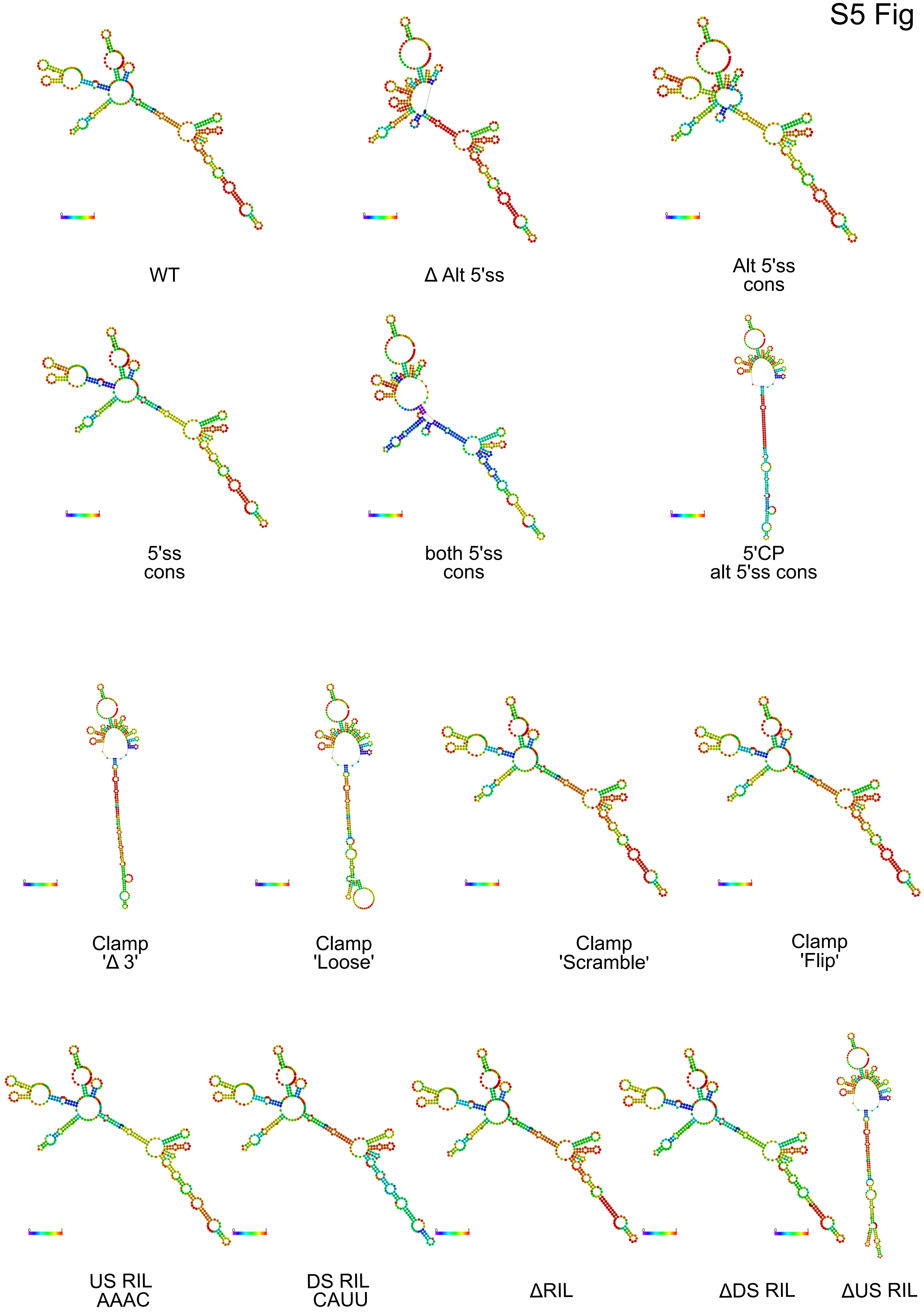

WT
Δ Alt 5'ss
Alt 5'ss cons
5'ss cons
both 5'ss cons
5'CP alt 5'ss cons
Clamp 'Δ 3'
Clamp 'Loose'
Clamp 'Scramble'
Clamp 'Flip'
US RIL AAAC
DS RIL CAUU
ΔRIL
ΔDS RIL
ΔUS RIL

S6 Fig

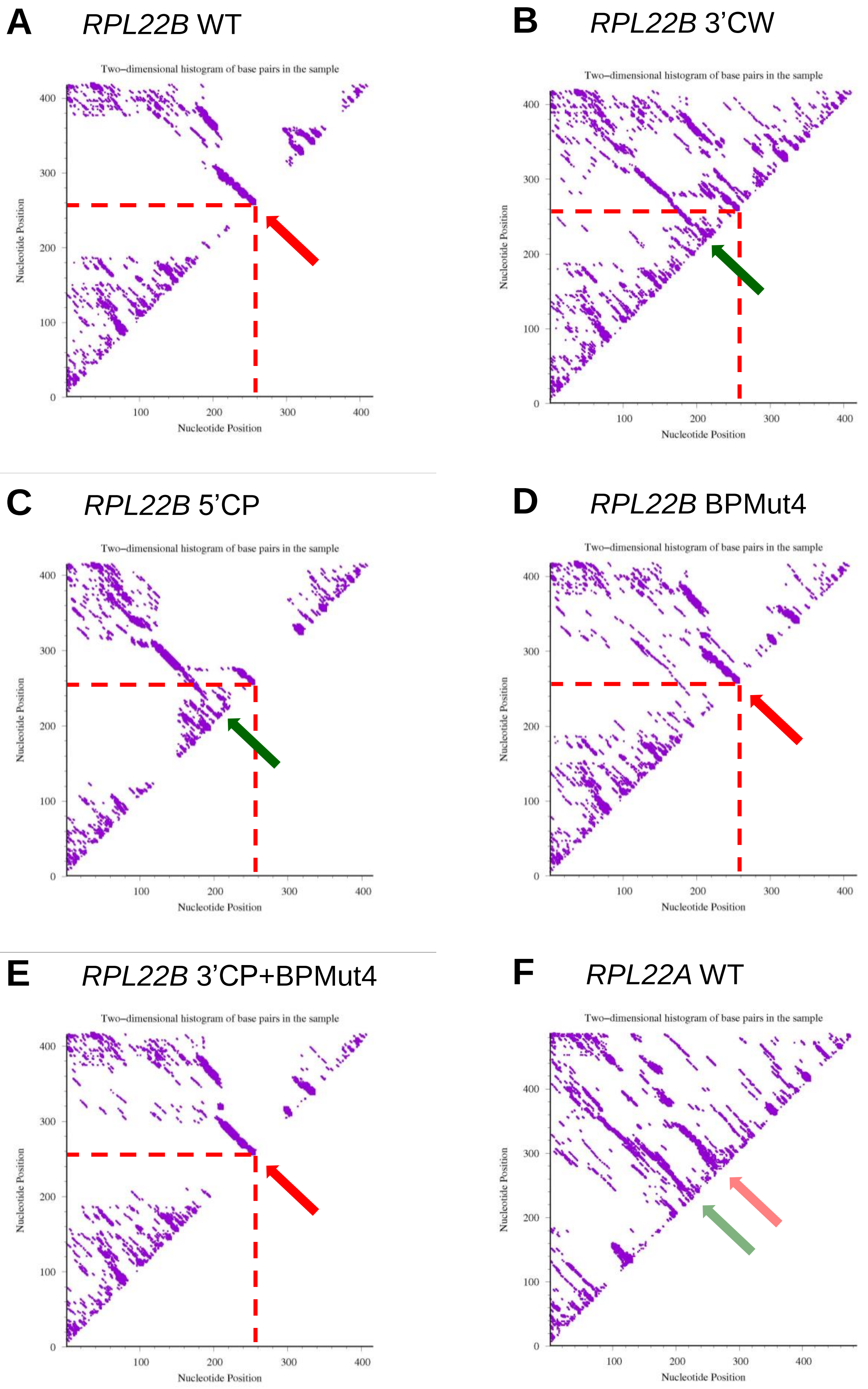
A
*RPL22B* WT
B
*RPL22B* 3'CW
C
*RPL22B* 5'CP
D
*RPL22B* BPMut4
E
*RPL22B* 3'CP+BPMut4
F
*RPL22A* WT
Two–dimensional histogram of base pairs in the sample
Nucleotide Position
Nucleotide Position
0
100
200
300
400

S7 Fig

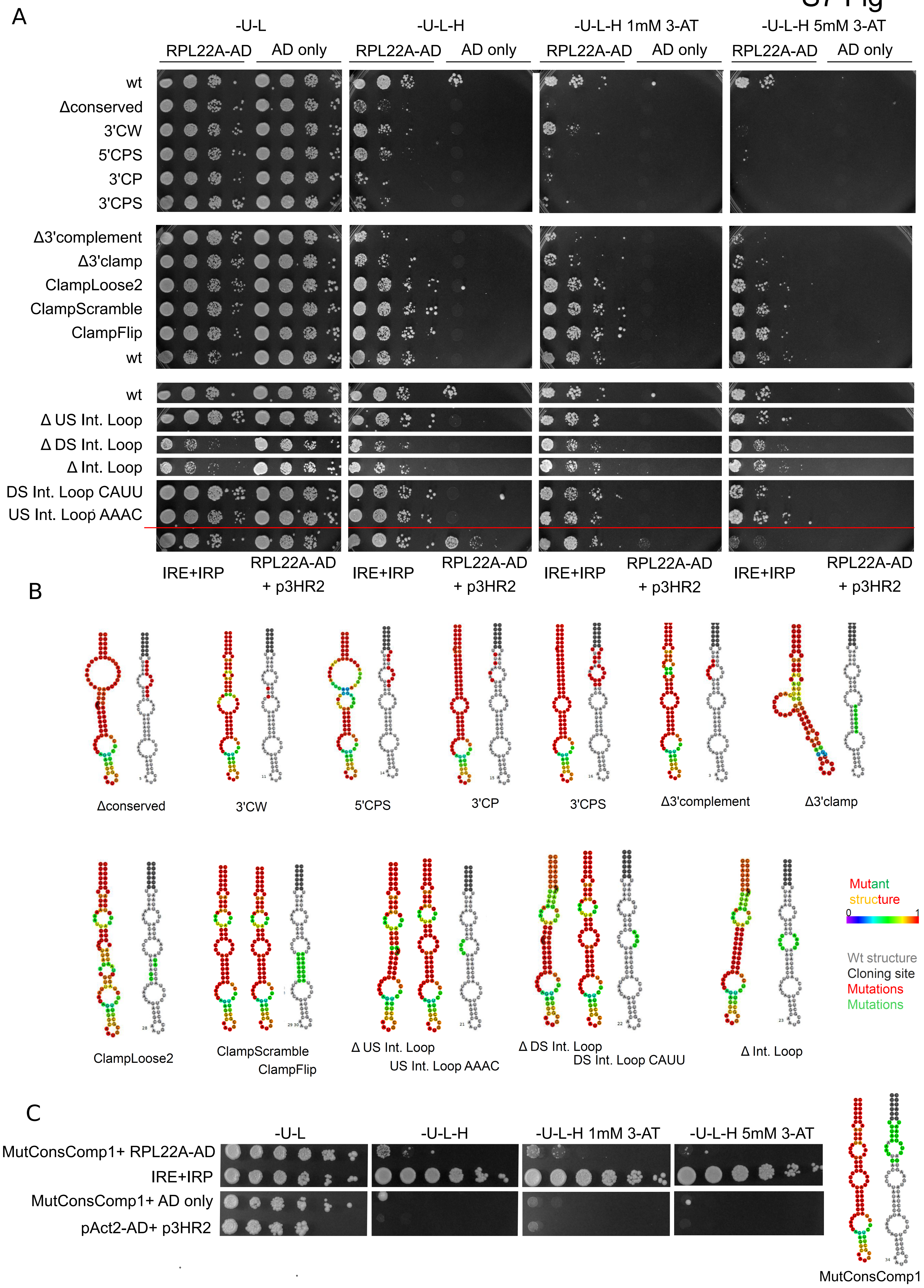
A
-U-L
-U-L-H
-U-L-H 1mM 3-AT
-U-L-H 5mM 3-AT
RPL22A-AD
AD only
wt
Δconserved
3'CW
5'CPS
3'CP
3'CPS
Δ3'complement
Δ3'clamp
ClampLoose2
ClampScramble
ClampFlip
wt
wt
Δ US Int. Loop
Δ DS Int. Loop
Δ Int. Loop
DS Int. Loop CAUU
US Int. Loop AAAC
IRE+IRP
RPL22A-AD + p3HR2
B
Δconserved
3'CW
5'CPS
3'CP
3'CPS
Δ3'complement
Δ3'clamp
ClampLoose2
ClampScramble
ClampFlip
Δ US Int. Loop
US Int. Loop AAAC
Δ DS Int. Loop
DS Int. Loop CAUU
Δ Int. Loop
Mutant structure
0
1
Wt structure
Cloning site
Mutations
Mutations
C
-U-L
-U-L-H
-U-L-H 1mM 3-AT
-U-L-H 5mM 3-AT
MutConsComp1+ RPL22A-AD
IRE+IRP
MutConsComp1+ AD only
pAct2-AD+ p3HR2
MutConsComp1

S8 Fig

```
Rpl22A    MAPN------TSRKQKIAKTFTVDVSSPTENGVFDPASYAKYLIDHIKVEGAVGNLGNAVTVTEDGTV   62
Rpl22B    MAPN------TSRKQKVIKTLTVDVSSPTENGVFDPASYSKYLIDHIKVDGAVGNLGNAIEVTEDGSI   62
KlRpl22   MAPN------TARKQKITKTFTVDVSSPTENGVFDPASYAKYLIDHIKVEGHVGNLGQAITVEEDGSV   62
CgRpl22   MAPN------TARKQKVVKTFTVDVSAPTENGVFDPASYAKYLIDHIKVENVVGNLGNAITVEEDGSV   62
CaRpl22   MAPV------TSKKSKSVKKFVVDVAAPVENDVFDQESYVKYLVEHVKVDGIVGNLGNDISITAESDN   62
SpRpl22   ---------MVKKNTKVSNKYIIDATAAVNDKIFDVAAFEKYLIDRIKVDGKTGNLGSSVSITEEGNK   59
DhRpl22   MAPI------TTKKNTAAKKLVVDTSAPTENGVFDQESYVKFLIENIKVEGIPGNLGNSIVTIERSKS   62
HsRpl22   MAPVKKLVVKGGKKKKQVLKFTLDCTHPVEDGIMDAANFEQFLQERIKVNGKAGNLGGGVVVSREGSS   68
YlRpl22   MAPV------KTQ---KANKYTVDCKAPSADGIFDVSSFEKFLTERIKVEGRTNQLGEDIKVSSNGDI   59
                      :      .  :*      : ::*   : ::* :.:**:.  .:**  :     .

Rpl22A    V---TVVSTAKFSGKYLKYLTKKYLKKNQLRDWIRFVSTKTNEYRLAFYQVTPEEDEEEDEE-   121
Rpl22B    V---TVVSSAKFSGKYLKYLTKKYLKKNQLRDWIRFVSIRQNQYKLVFYQVTPEDADEEEDDE   122
KlRpl22   V---TIVSTTKFSGKYLKYLTKKYLKKNQLRDWIRFVSTKTNEYKLAFYQITPEDEEEEEDEE   122
CgRpl22   V---TIVSTTKFSGKYLKYLTKKYLKKNQLRDWIRFVSTKTNSYRLAFYQVTPEEEEEDEE--   120
CaRpl22   KVVVVVVSGNGSFSGKYLKYLTKKYLKKNQIRDWIRFVSVKQNQYKLQFYAVAEDDEEEEDEE-  124
SpRpl22   K--IAVIAHIDFSGRYLKYLTKKFLKKHSLRDWLRVVSTKKGVYELRYYNVVVGNDEEEQ---   117
DhRpl22   V---VVVSNTKFSGKYLKYLTKRYLKKNQIRDWIRFVSVKQNQYQLQFYSVADDEEEEDEE--   120
HsRpl22   K--ITVTSEVPFSKRYLKYLTKKYLKKNNLRDWLRVVANSKESYELRYFQINQDEEEEEDED-   128
YlRpl22   V---TVVSTTQFSGKYLKYLTKKYLKKQQLRDWIRVISTSKGNYTLKFYNVVANEEDEE----   115

              .: .   ** :*******::***:.:***:*.::     * * :: :   : :*:
```

S9 Fig

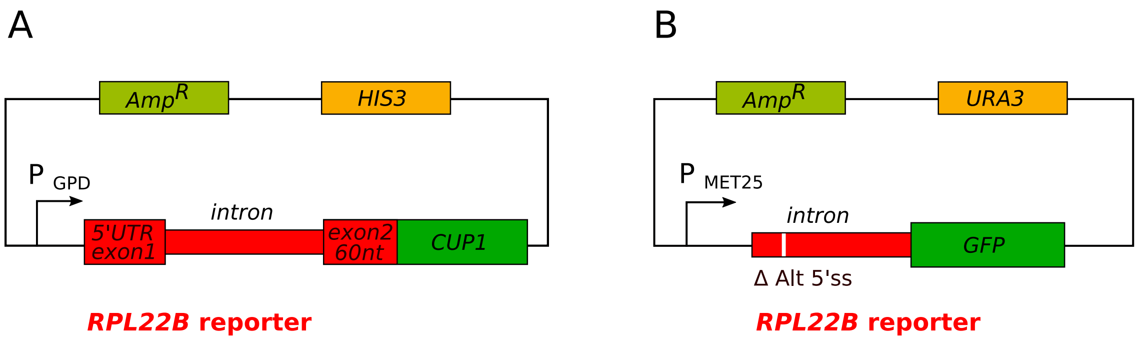
A
AmpR
HIS3
P GPD
intron
5'UTR exon1
exon2 60nt
CUP1
RPL22B reporter
B
AmpR
URA3
P MET25
intron
GFP
Δ Alt 5'ss
RPL22B reporter

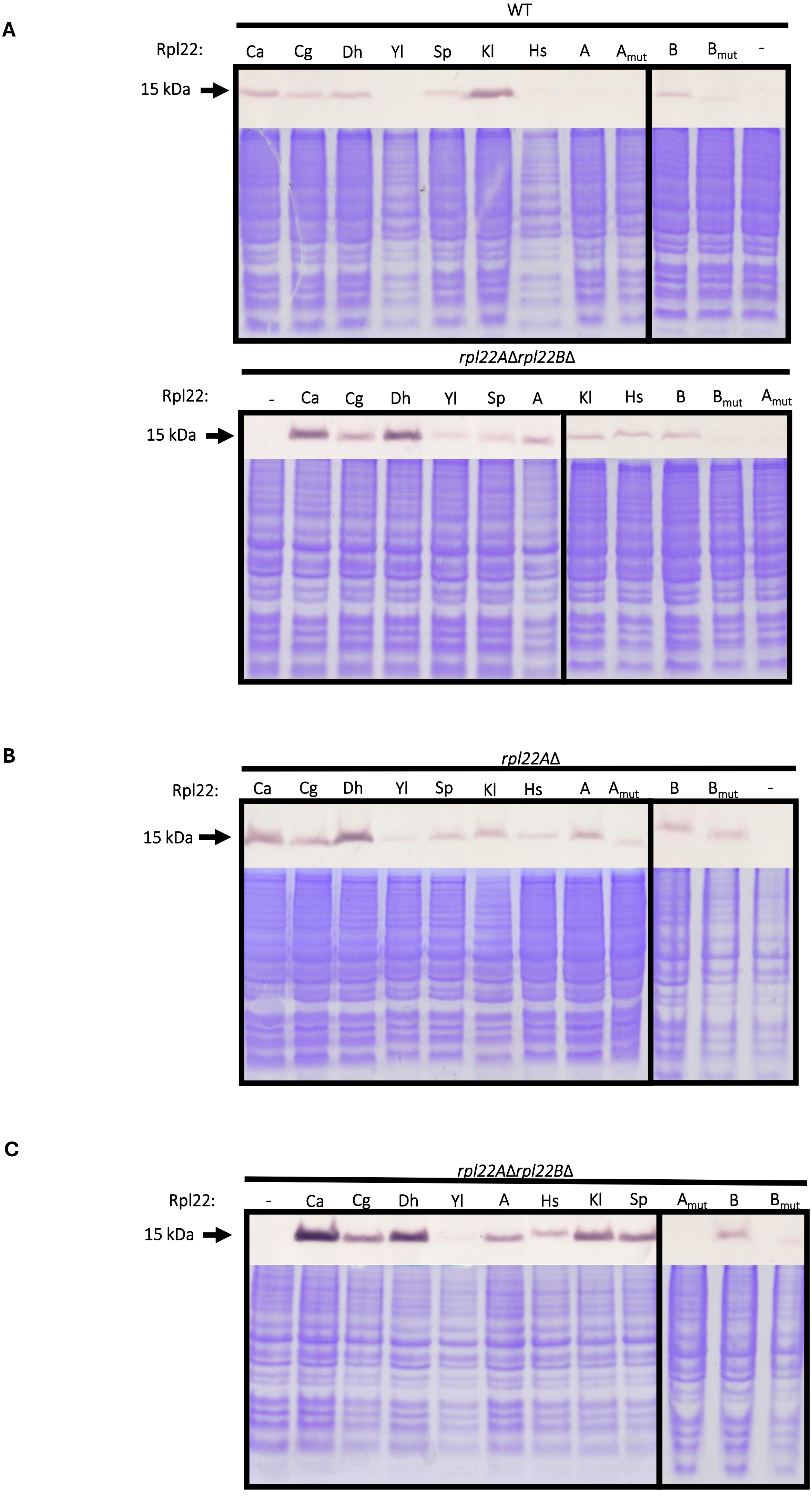
A
WT
Rpl22: Ca Cg Dh Yl Sp Kl Hs A A$_{mut}$ B B$_{mut}$ -
15 kDa
rpl22AΔrpl22BΔ
Rpl22: - Ca Cg Dh Yl Sp A Kl Hs B B$_{mut}$ A$_{mut}$
15 kDa
B
rpl22AΔ
Rpl22: Ca Cg Dh Yl Sp Kl Hs A A$_{mut}$ B B$_{mut}$ -
15 kDa
C
rpl22AΔrpl22BΔ
Rpl22: - Ca Cg Dh Yl A Hs Kl Sp A$_{mut}$ B B$_{mut}$
15 kDa

S10 Fig
